# Climate Change Controls Phosphorus Transport from the Watershed into Lake Kinneret (Israel)

**DOI:** 10.1101/2021.03.16.435628

**Authors:** Moshe Gophen

## Abstract

Part of the Kinneret watershed, the Hula Valley, was modified from wetlands – shallow lake for agricultural cultivation. Enhancement of nutrient fluxes into Lake Kinneret was predicted. Therefore, a reclamation project was implemented and eco-tourism partly replaced agriculture. Since the mid-1980s, regional climate change has been documented. Statistical evaluation of long-term records of TP (Total Phosphorus) concentrations in headwaters and potential resources in the Hula Valley was carried out to identify efficient management design targets. Significant correlation between major headwater river discharge and TP concentration was indicated, whilst the impact of external fertilizer loads and 50,000 winter migratory cranes was probably negligible. Nevertheless, confirmed severe bdamage to agricultural crops carried out by cranes led to their maximal deportation and optimization of their feeding policy. Consequently, the continuation of the present management is recommended.

## 1) Introduction

Phosphorus, manganese, and organic matter relations in the Hula Valley peat soil have been widely studied (Gophen et al 2014; Haygarth et al. 2013; Litaor et al. 2013; 2014; Reichman et al. 2013; 2016; Yatom and Rabinovich 1999; Yatom et al.1996;Xiang et al 1996; Barnea 2009). The impact of geochemical parameters of pH, oxidation–reduction (redox), wettability (rainfall, irrigation), soil properties, temperature, and agricultural management (fertilization) conditions on the dynamics of phosphorus and peat soil particle bound–release relations had been thoroughly investigated in those studies. Moreover, plant-mediated phosphorus and its role in the Phosphorus transport from altered wetland soils into water pathways has been documented as well (Gophen 2000; Simhayov et al. 2013). External sources of phosphorus in dust deposition and agricultural fertilization have been studied (Foner et al. 2009; Litaor et al. 2013; Barnea 2009; Reichman et al 2013). Phosphorus supply into the Hula ecosystem by the winter migrators cranes (Grus grus) fed by corn seeds has also been documented (Gophen 2017). Information on phosphorus transportation and migration in relation to topography, hydrology, vegetation coverage, and land use management has been widely discussed (Reddy et al. 1999). Hydrological linkage between the Hula Valley and the downstream Lake Kinneret makes the dynamics of water-mediated phosphorus input into the lake essential. The scope of this paper includes the long-term (1970-2018) search for the bound between phosphorus resources in the Hula Valley and its water-mediated transport into Lake Kinneret. In other words, what is the fate of phosphorus sources in the Hula Valley? Does land-use-and land-cover management policy obtainable in the Hula Valley control the Valley’s entire stock of phosphorus? Does water-mediated phosphorus effluents include total phosphorus capacity, and if not, what is the fate of the rest?

### 1.2.) Regional Hydrology

The Hula Valley and Lake Kinneret are located in the Syrian–African Rift Valley in northern Israel (Figure 1). Lake Kinneret is the only natural freshwater lake in Israel. Until 2010, an average of 336 mcm (336 million cubic meters) were pumped annually (34% in winter and 66% in summer) from the lake and supplied mostly for domestic usage and partly for agricultural irrigation (Gvirtzman 2002). Since 2010, desalinization plants have supplied almost the full water demand for domestic consumption, reducing dramatically the pumping rate from Kinneret. The water quality of Lake Kinneret is a national concern since pollutant (TP included) inputs from the Hula Valley are prominent. Over 95% of Israel’s natural water resources are utilized. The total national water supply is 2.11 bcm (2.11 billion cubic meters), of which 0.55 bcm comes from the Kinneret–Jordan water system and 0.7 bcm from desalinization. The area of the Kinneret drainage basin is 2730 km^2^; it is located mostly to the north of the lake from which the Hula Valley is about 200 km^2^. Three major headwater rivers (Hatzbani, Banyas and Dan) flow from the Hermon Mountain region (Figure 1) located in the northern part of the Kinneret drainage basin (2730 km^2^). These rivers join the River Jordan which, before the Hula drainage,operation crosses the Valley through two branches (tributaries) flowing into the old Lake Hula. From the Lake Hula at an altitude of about 61 mamsl (61 meters above mean sea level), the River Jordan flows downstream into Lake Kinneret at an altitude of 209 mbmsl (209 meters below mean sea level) for a distance of about 15 km. The Jordan River contributes about 63% of the Kinneret water budget and more than 50% of the total external nutrient inputs (Gvirtzman 2002). Before the drainage of Hula Valley (1957), the land was covered by Lake Hula (1.5 m mean depth; 13 km^2^ water surface) and 3500 ha of swamps. The swampy area was completely covered by water in winter and partly covered in summer. To the north of the swamps was an area (3200 ha) where water table levels were high in winter, making agricultural cultivation impossible. During summer periods, when underground water levels declined, this 3200-ha surface was successfully cultivated.

**Figure 1:**
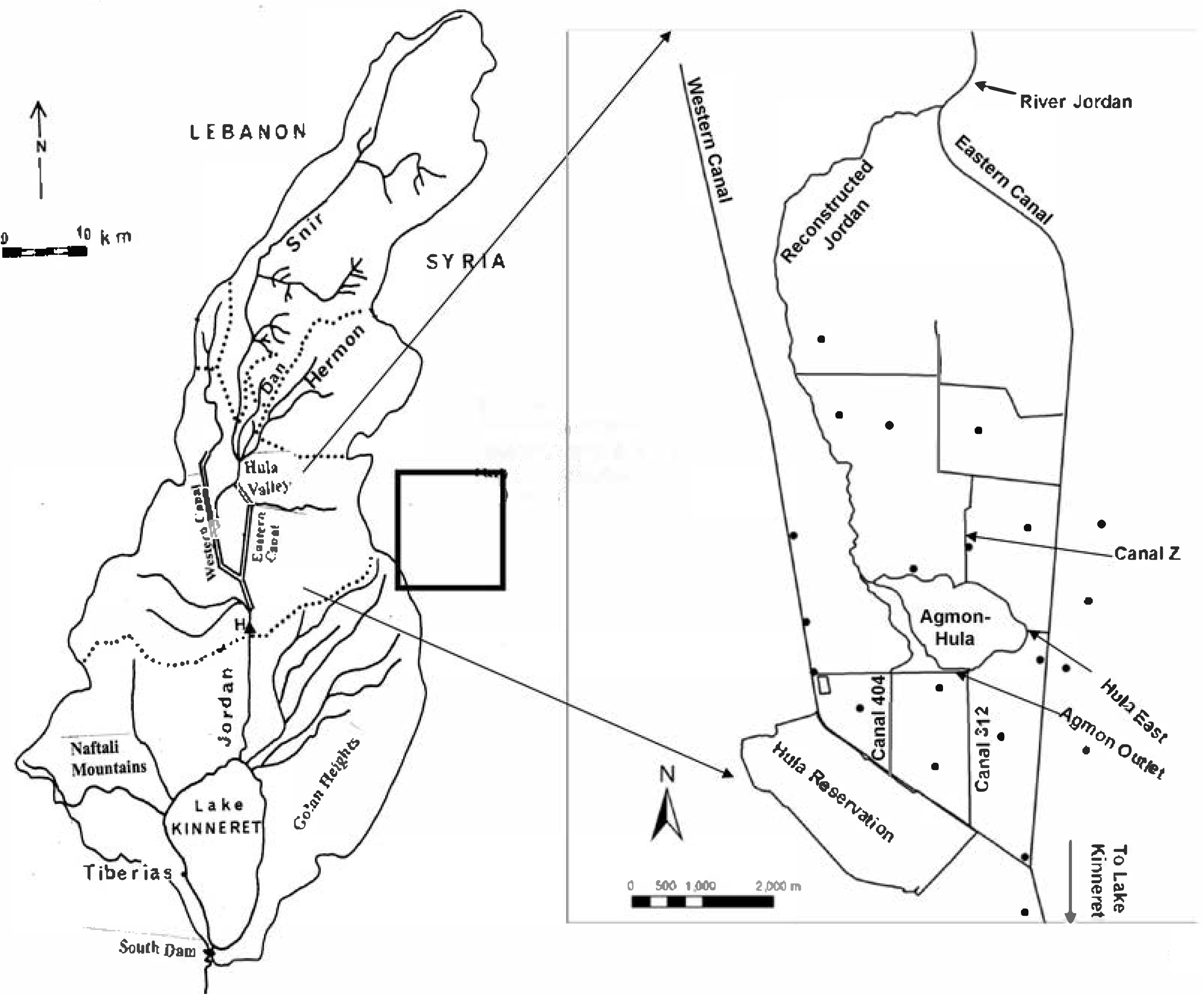
Geographical Maps: Kinneret watershed (left) Hula Valley (right) Drills (⚫) are indicated

### 1.3.) Anthropocene History of the Hula Valley (Karmon 1956)

The Hula Valley was turned into a wilderness by the Mongolian from 1240 AC. Mosquitos carrying malaria were introduced into the Hula Valley by the Crusaders, and the inundation of the Hula Valley was enhanced as a result of the construction of the Benot Yaakov bridge downstream by Bivers in 1260. Later on, malaria became a major parameter which affected human activity in the Hula wetlands.

The Ghawarna tribes were the first people to settle in the Hula Valley in the 4^th^ decade of the 19^th^ century but their settlement came to an abrupt end in 1948.Settling in the Hula Valley by the Ghawarna tribes was initiated during the 4^th^ decade of the 19^th^ century and came to an end abruptly in 1948. The development of the Ghawarna settlement was very slow during the 19^th^ century but significantly accelerated during the 1st half of the 20^th^ century. According to British sources, between 1877 and 1948, the Ghawarna population increased from 520 to 31,470. Prior to 1830, there were no permanent settlements of the Ghawarna in the Hula Valley. Residents from the northern region came down from the surrounding mountains with their cattle herds for grazing and for agricultural cropping in summer and stayed most of the summer months in the parts of the valley that were not inundated. This was the summer paradox of the Hula wetlands prior to the 20^th^ century: the drier the winter, the more the quantity of grass for cattle grazing and land for agriculture. The cultivated land was like a puzzle of plots, namely “Mazraa” (or “Azeva”).

The history of Jewish settlements in the upper Galilee and particularly in the Hula Valley and its vicinity dates back to the end of the 19^th^ century and the beginning of the 20^th^ century. Nevertheless, extensive Jewish settlement in the Hula Valley region started in the 1940s. The building of the drainage of the old Lake Hula and adjacent swampy area began in 1950 and was completed in 1955. The Old Lake Hula wetland area was converted to arable land. Beneficial crops were produced but not without difficulties.

### 1.4.) The Hula Reclamation Project

Because of the Lake & wetland drainage (1957), more than 6500 ha of natural wetland area were converted for agricultural development. Therefore, the unique natural composition of fauna and flora of exceptional diversity was almost demolished. The newly created arable land became a source of income to the residents of northern Israel. For 40 years it was successfully cultivated and agricultural products (mostly cotton, corn, alfalfa, and vegetables) were economically produced, and nutrient flux into Lake Kinneret did not threaten the lake’s water quality. Nevertheless, as a result of inappropriate management, drainage canals were blocked, irrigation methods were not suitable for optimal soil management and fertility, and crop utilization and water tables declined. Consequently, the soil structure of the upper layers (0–0.5m) became oxidized and deteriorated, heavy dust storms became frequent, and the soil surface subsided (7–10 cm/year). Due to the decline in the water table level and longer periods of leaving bare and dry soils uncultivated, underground fires occurred quite often. increased Rodent population outbreaks caused severe damage to agricultural crops and the stability of drainage canal banks. In the 1980s intensive cultivation of the land was gradually abandoned. Therefore, in the period 1990–1997, the whole drainage area went through a reclamation project, referred as the Hula Project, which was focused on the 500 ha in the middle part of the valley at the lowest altitude. The project was aimed primarily at reduction of nutrient fluxes from the Hula Valley soil while implementing modern irrigation methods to reintroduce economical land use and integrate eco-tourism. The reclamation project included several operational elements, viz.: increasing the soil moisture by elevating the ground water table (GWT), changing the irrigation method and renewing the drainage system in the entire valley, and creating a new shallow lake called Agmon-Hula. The surface area and mean depth of this lake were 110 ha and 0.5 m respectively in the years 1994-2010 but later these value became to 82 ha and 0.2 m. (Gophen et al. 2003). This shallow lake was designed to be operated as a drainage basin for the valley and provide an ecological service of eco-touristic wetland. A plastic sheet (4-mm thickness) was placed vertically (0–4.5 m) over a distance of 2.8 km, crossing the east-west direction, the west-southern part of the valley, to separate the ground water tables and to prevent underground migrated leakageg of pollutants downstream to Lake Kinneret.

Prior to the drainage of the Hula Swamps and the Old Lake becoming a national concern, the interest in the north was security combined with demography and population dispersal accompanied by agricultural income sources. Later, the search for essential utilization of the Hula land became a national concern. Optimized implementations of agricultural technologies were not easily established and plenty of difficulties interrupted their efficient utilization. The Hula Project included the development of a new multipurpose shallow lake known as “Agmon-Hula” (Gophen et al. 2016). The objective of this new lake was the creation of a sufficient hydrological volume to collect peat soil-drained, nutrient-rich water effluents mixed with fresh Jordan River waters to prevent deterioration of water quality. Nutrient-rich polluted waters from Lake Agmon-Hula were transferred for irrigation usage outside the Kinneret drainage basin. Agmon-Hula and the surroundings (500 ha) were earmarked for commercial eco-touristic management. Natural attractions were designed for observational touring of aquatic vegetation landscape, bird watching and sport fishing recreation. The rationale was to replace agriculture with another income source for the land owners. The original design was successfully implemented and crane wintering provided an attractive experience for tourists.

The objective of this paper is to get an insight on phosphorus resources in the Kinneret watershed ecosystem and to evaluate the practical contributions of phosphorus to this ecosystem.

## 2.) Material and Methods

### 2.1.) Data Sources

Ground water table, total phosphorus (TP) concentrations in the water canals of the Hula Project area and Agmon-Hula effluent (1994–2020), the discharges of the three headwater rivers and River Jordan discharges (1970–2018) (mcm/y; 10^6^ m^3^) and rainfall (1940–2020) were statistically evaluated. Maximal counts of wintering cranes in the period 1997–2020 were also evaluated. Data were obtained from the following sources: Annual Reports of the Kinneret Limnological Laboratory (LKDB-IOLR 1970-2018); National Meteorological Service; National Hydrological Service; National Water Authority; MIGAL-Scientific Research Institute; Mekorot Water Supply Company Ltd. (Nazareth, Israel); Monitoring Unit Jordan District. Data on the Jordan River nutrient loads, concentrations, and discharge were obtained from the annual and temporal reports. Agmon-Hula TP concentrations (1993–2019) were obtained from the annual reports (Gophen and D. Levanon 1993–2006; Gonen 2007; Barnea 2008, 2008–2018). Monthly means (1993–2019) of 277 sampled TP concentrations of Agmon-Hula effluents were evaluated; twenty-seven months (9 in 2016) were not sampled in this period due to technical difficulties. Monthly averages of 2–6 weekly samples of TP concentrations in underground water samples collected monthly (14) from the top level of the ground water table (GWT) in 14 drills distributed in the Hula valley were reconsidered as well (Figure 1) (Gophen et al. 2014).

### 2.2.) Statistical Methods

Statistical analyses (fractional polynomial regression) (FP) were carried out using STATA 9.1, Statistics-Data Analysis, Chapter fracpoly-Fractional Polynomial regression; StataCorp, 2005, Stata Statistical Software: Release 9. College Station, TX, USA: StataCorp LP. pp. 357–370 (See also: Royston, P. and D. G. Altman, 1994). Regression was carried out using fractional polynomial of continuous covariates: Parsimonious parametric modeling (with discussion), Applied Statistics 43: 429–467. The purpose of FPs is to increase the flexibility of the family of conventional polynomial models. Although polynomials are popular in data analysis, linear and quadratic functions are severely limited in their range of curve shapes, and cubic and higher-order curves often produce undesirable artifacts such as “edge effects” and “waves” (STATA 9).

## 3.) Results

Climate change conditions of precipitation decline and, consequently, Kinneret Headwater river discharges were indicated since the 1980s (Figure 2) (Gophen 2021; Reichman et al. 2016). Long-term fluctuations of the annual means of Total Phosphorus (TP), concentrations in the Agmon-Hula outflow, Jordan River (Figure 3 and 4) whilst slightly increased in the Kinneret epilimnion (Gophen 2021). The decline in TP concentration in Jordan River ranged between 0.21 and 0.14 ppm, and was accompanied by a decline in total nitrogen (TN) and total inorganic nitrogen (NORG) (Figure 5). The increase in TP concentration in the Kinneret epilimnion was from 0.015 to 0.021 ppm (Gophen 2021). Moreover, the TP concentration dynamics in the Agmon effluent and Jordan waters indicates an inverse relation (Figure 6). The trend of ground water table decline since 2010, which is due to the climatological dryness trait (Figure 7), probably influenced TP dynamics in the peat organic soil. Temporal (1994–2020) changes in TP concentrations in the Agmon-Hula effluent indicates a long-term trend of elevation (Figure 8,9). Nevertheless, the seasonal and annual dynamics of TP content in the Agmon-Hula waters (and obviously their outflow) constantly show significant elevation during late summer– autumn months, which is 6 months after the northern migration of cranes (Figure 10 and 11). The annual increase in TP in late summer–autumn is due to degradation and decomposition of submerged vegetation. Consequently, it is suggested that cranes do not contribute significantly to TP levels in Lake Kinneret, and the increase in the TP concentration of the epilimnion is the result of internal sources. Moreover, positive regressions (r^2^ = 0.596) were indicated between River Jordan discharge and nutrient inflow loads (p < 0.0001) for TP. Independently, the discharges in the Jordan River have declined since the mid-1980s from 15 to <10 m^3^/s, caused by precipitation decline.

**Figure 2:**
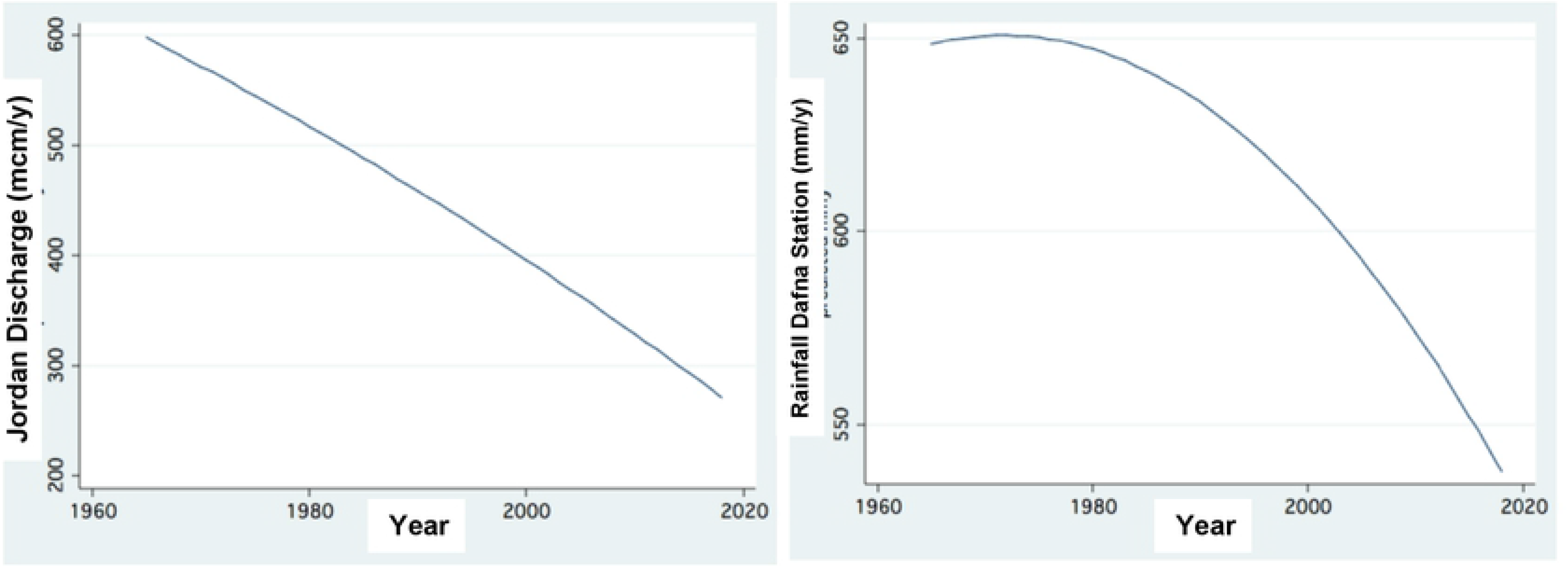
Climate Change in the Kinneret Region: Decline of Rainfall (mm/y) (right) accompanied by reduction of Jordan River Discharge (10^6m^3/y) (Left).

**Figure 3:**
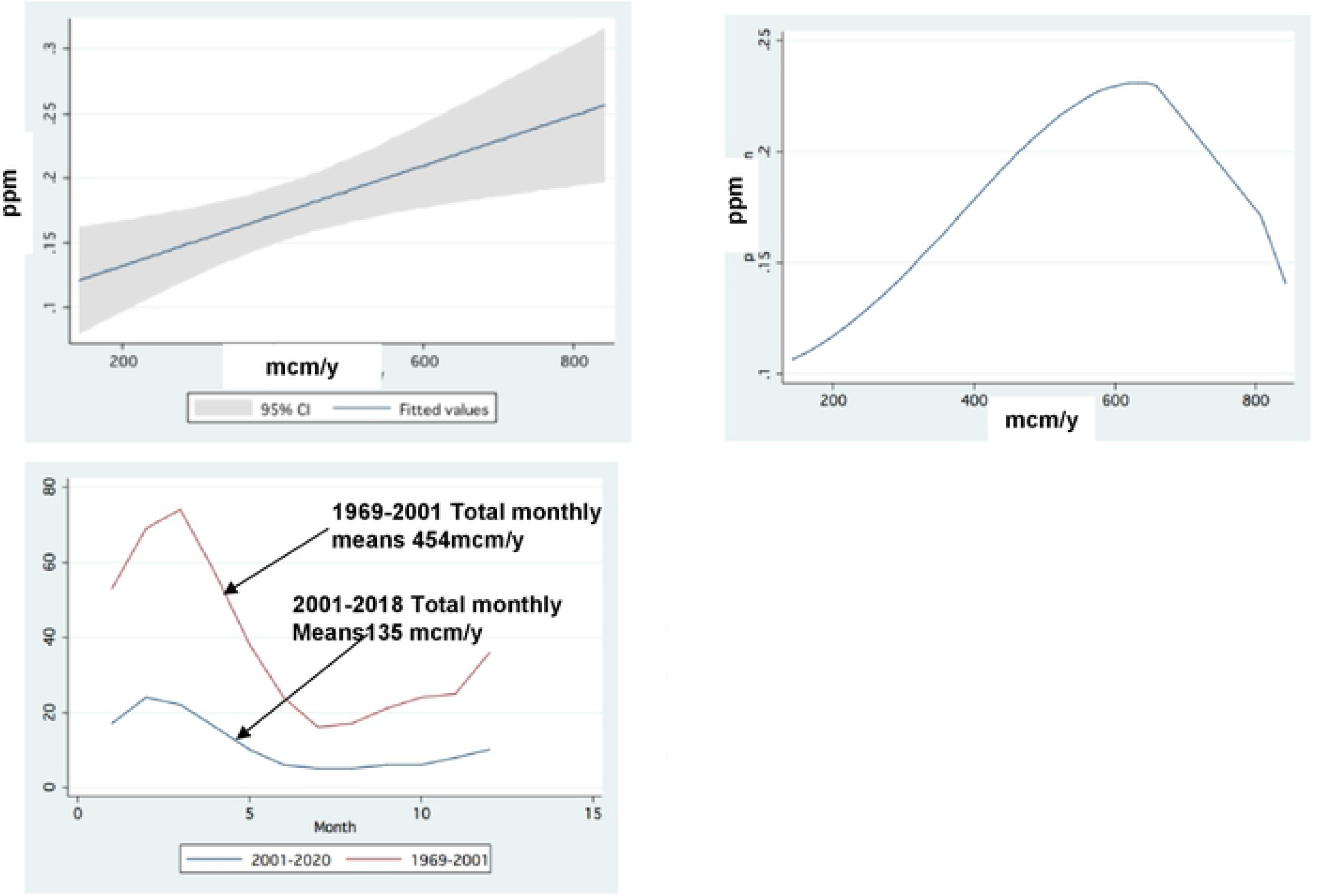
Upper Panels: Jordan Water Annual averages of TP concentration (ppm) Vs. Discharge (10^6 m^3/y): Left: Linear Prediction (95% CI); Right: Fractional Polynomial Regression. Lower panel: Monthly averages (1969-2001, and 2001-2018) of River Jordan discharge (mcm/m).

**Figure 4:**
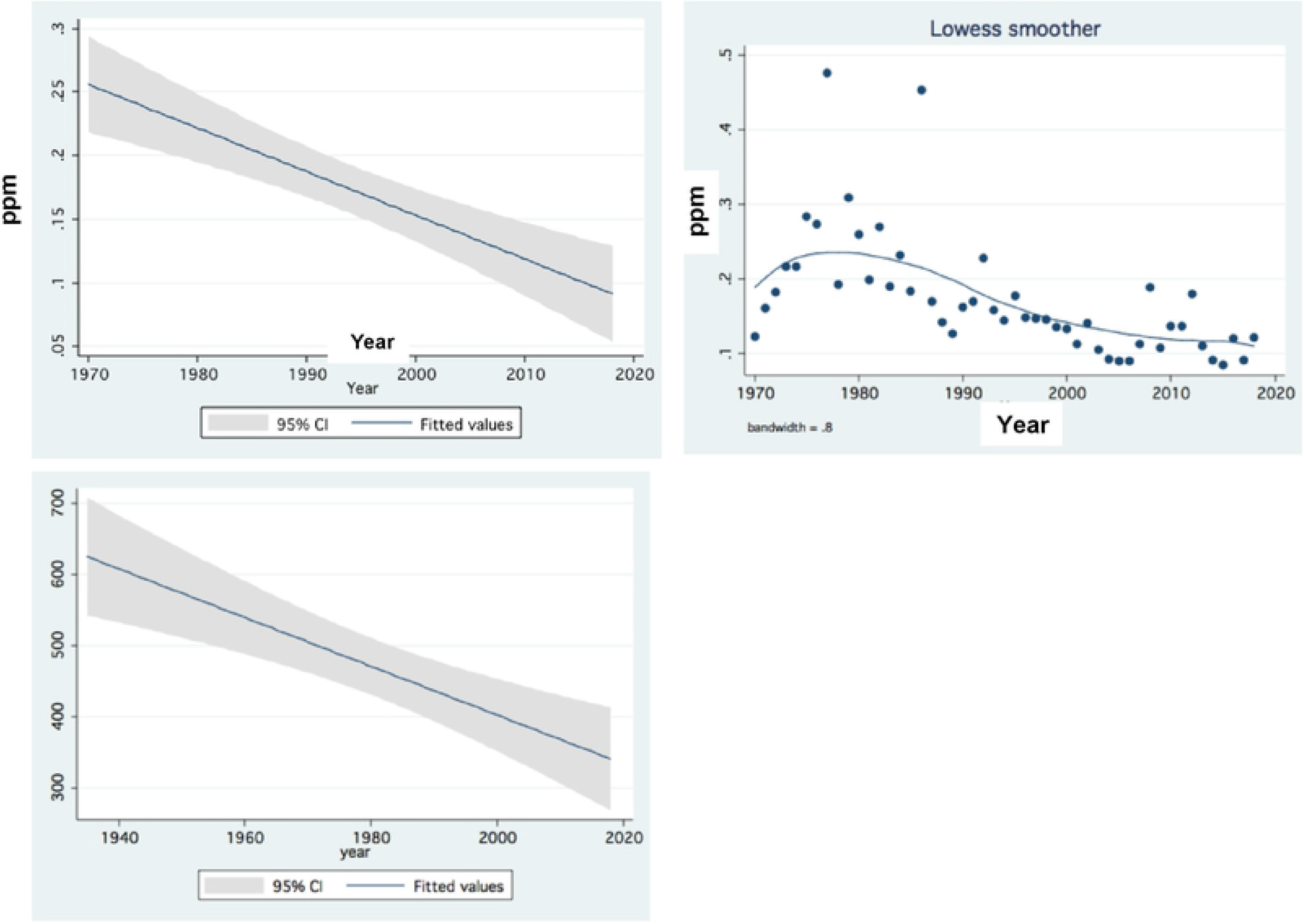
Temporal (1970-2018) changes of annual averages of TP concentration (ppm) in Jordan Water (upper panels): Left: Linear Prediction (95% CI) Right: Trend of Changes (LOWESS Smoother 0.8) Lower: Jordan annual Discharge (mcm/y (1940 - 2018).

**Figure 5:**
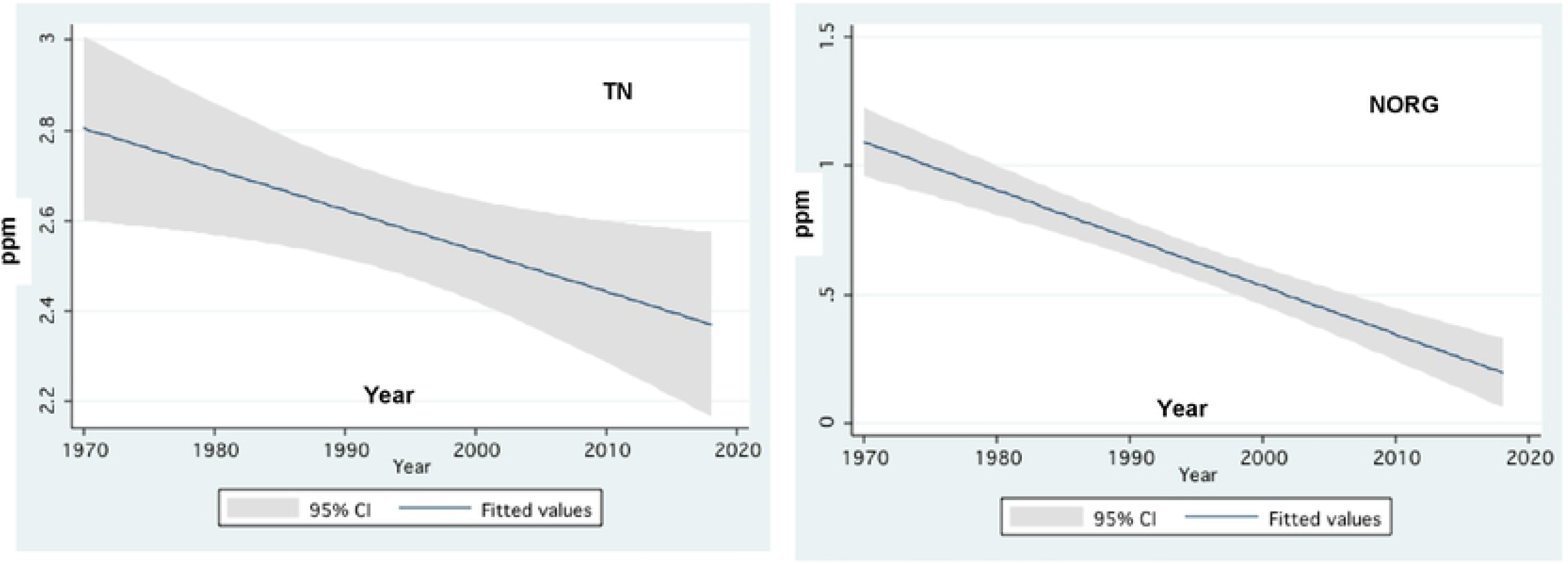
Linear Prediction (95% CI) of temporal changes of annual averages of Total Nitrogen (TN) (left) and Organic Nitrogen (NORG) (right) concentration (ppm) in the Jordan waters during 1970-2018.

**Figure 6:**
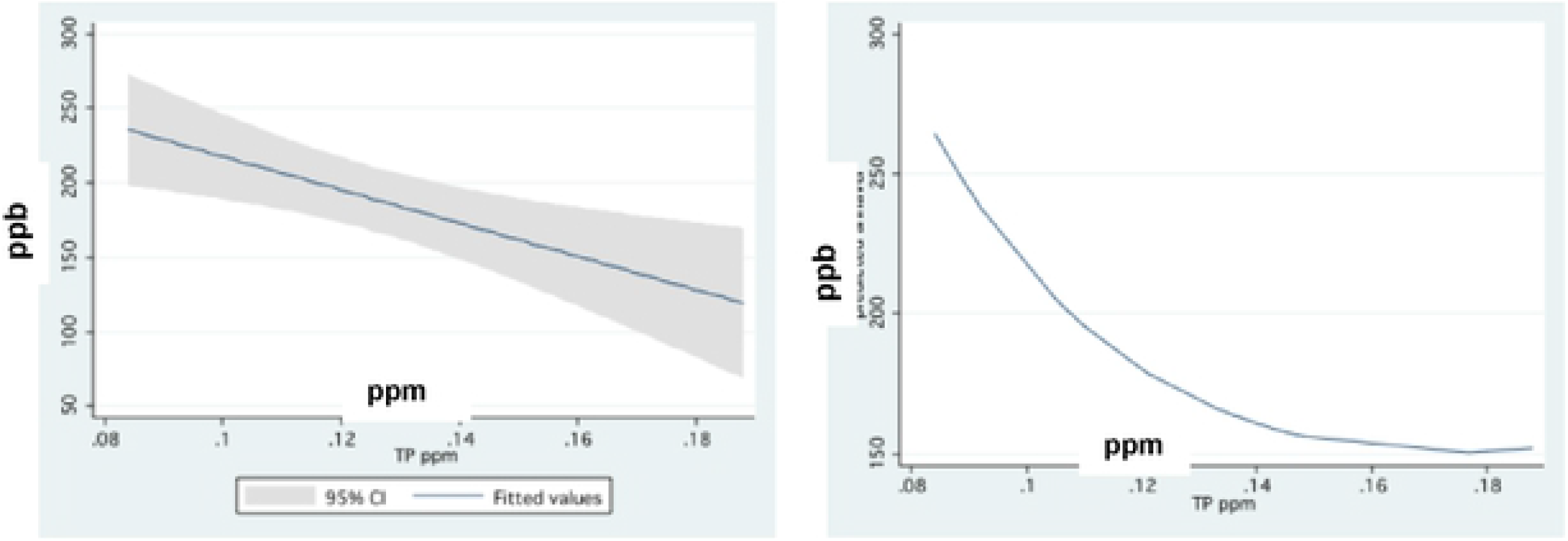
Annual Averages of Total Phosphorus (TP) concentrations (ppb) In the Agmon effluent Vs Annual Concentrations (ppnm) of TP in Jordan waters: Linear Prediction (95% CI) (left) and Fractional Polynomial Regression (right) during 1993-2018.

**Figure 7:**
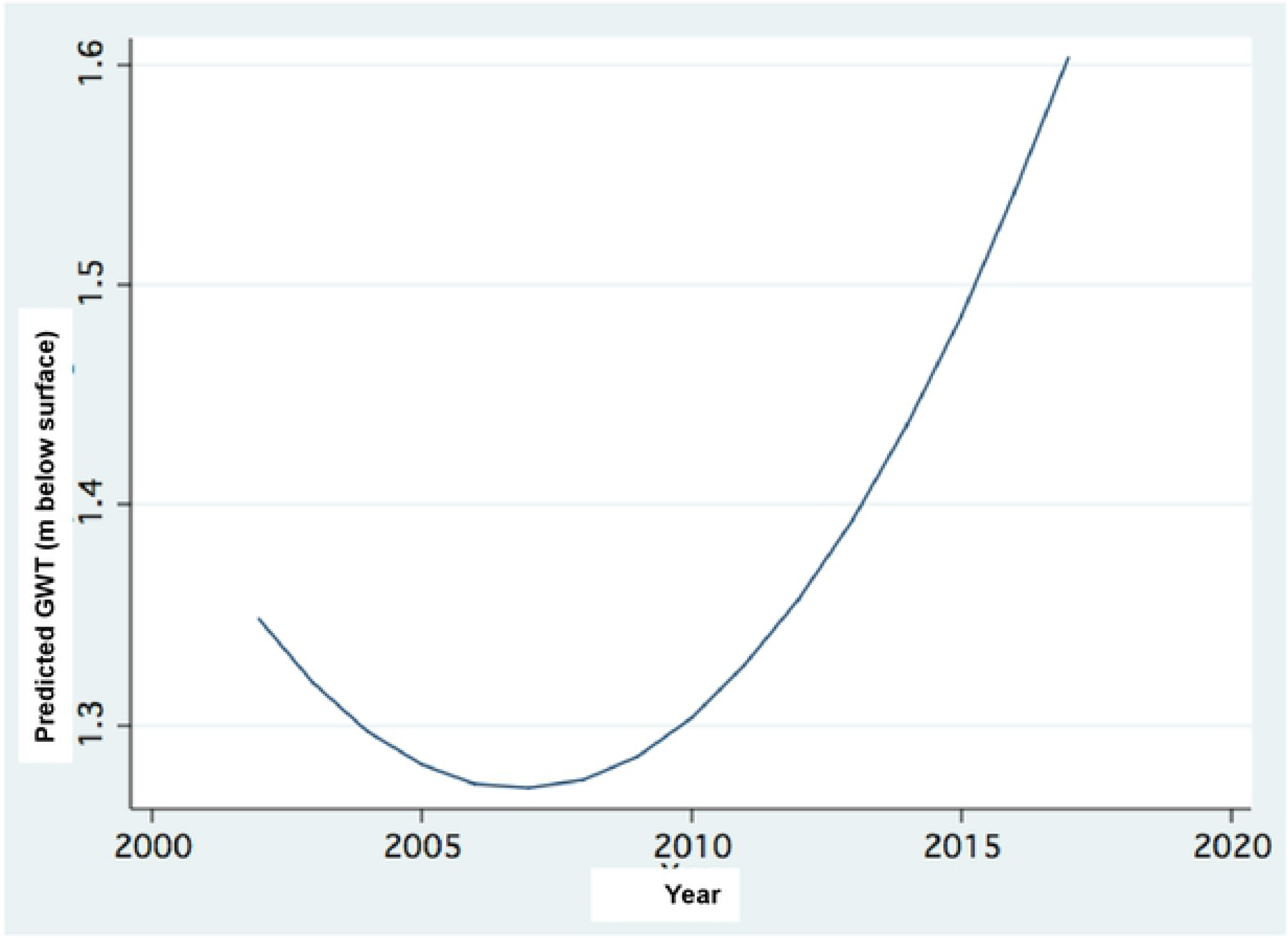
Climate change in the Kinneret region: 32 cm Decline of Ground Water Table (m below surface) in the Hula Valley: annual averages of bi-weekly measured in 32 drills distributed throughout the entire valley during 1993-2020.

**Figure 8:**
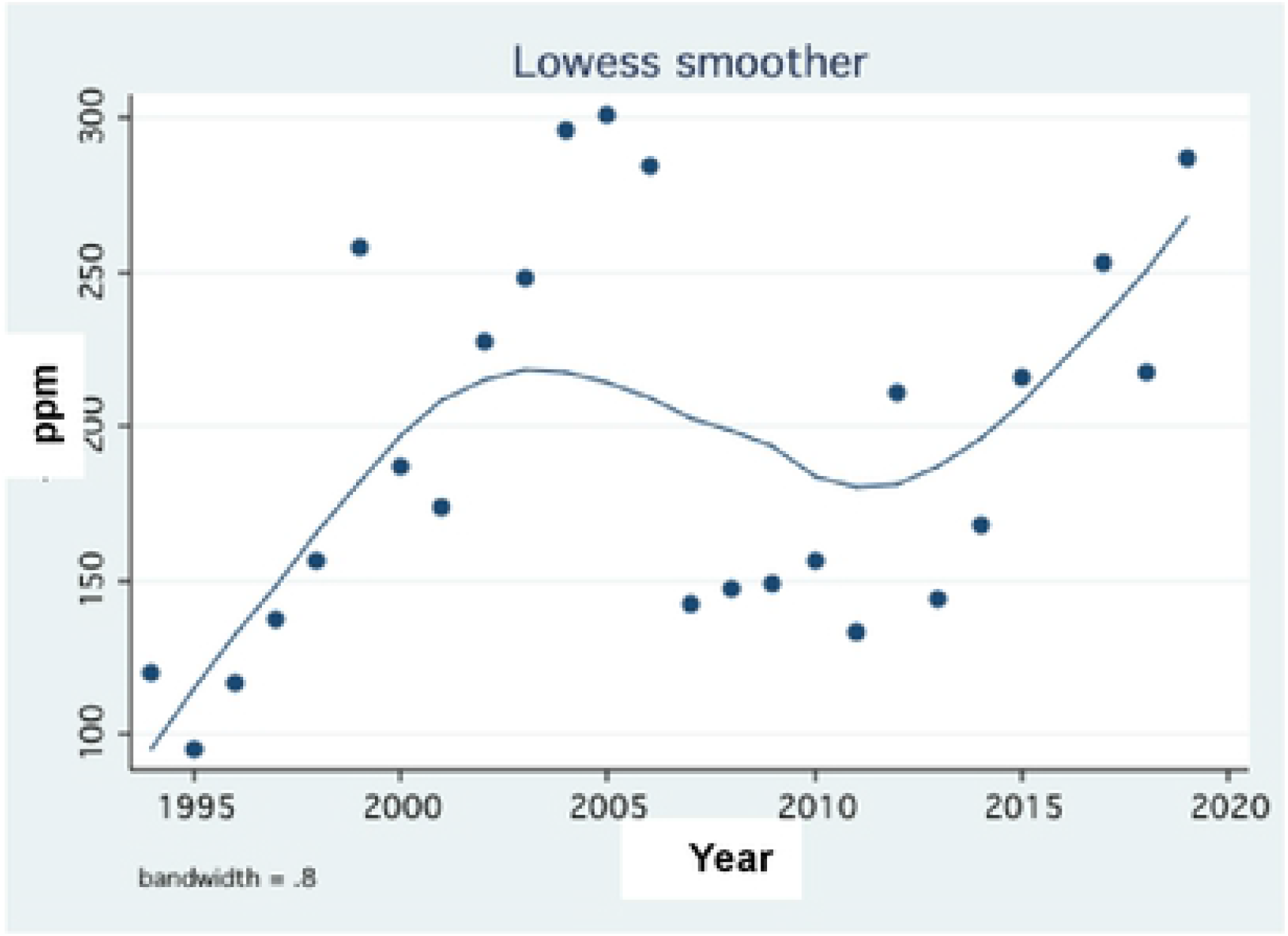
Lowess Smoother (Bandwidth 0.8) of Temporal changes of Total Phosphorus (TP) concentrations (ppb) in Lake Agmon-Hula Annual averages Vs. Years (1994-2020)

**Figure 9:**
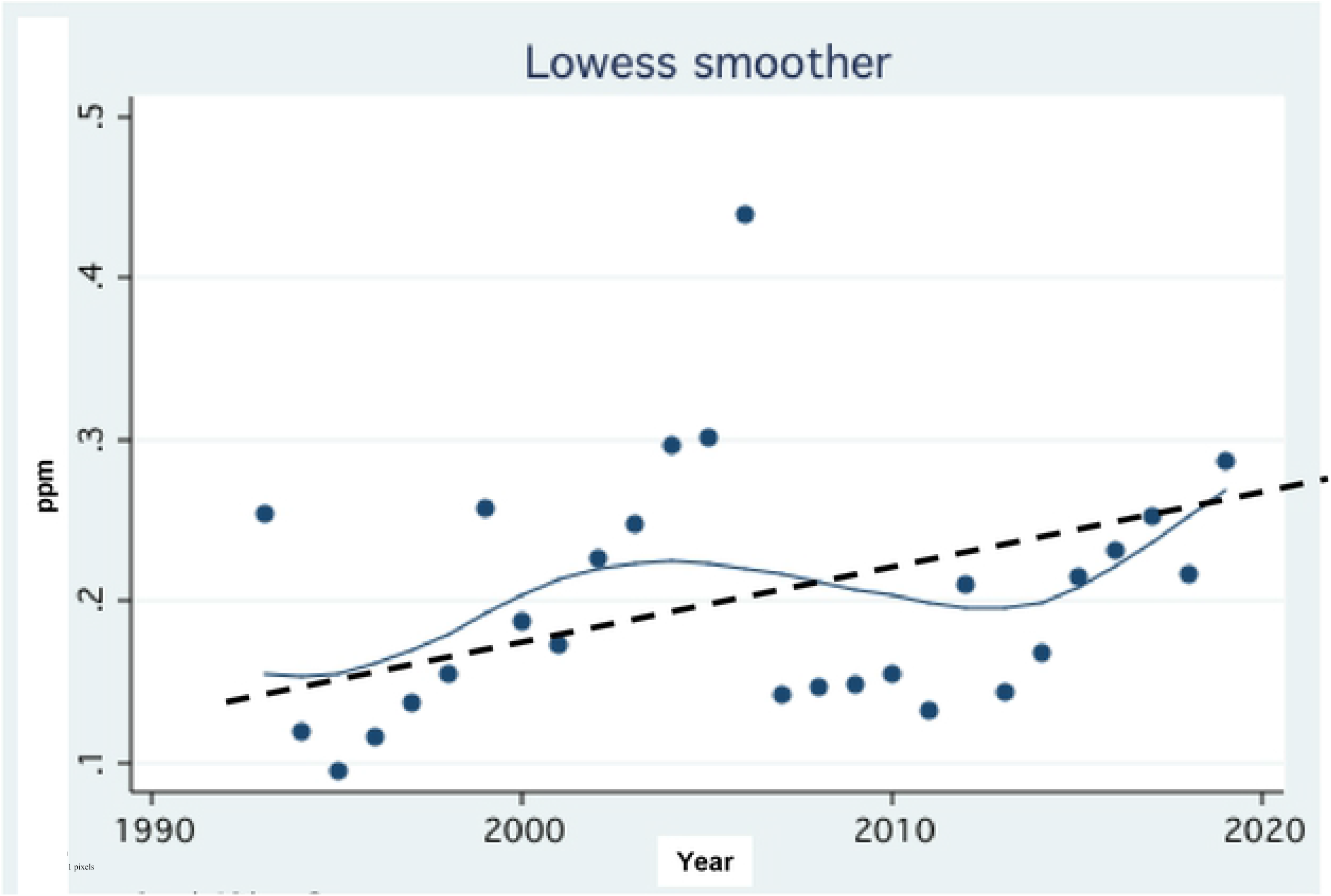
Trend of Changes (LOWQES Smoother 0.8) of Total Phosphorus concentration (TP; ppm) in the Lake Agmon-Hula effluent during 1993-2020; Avergaed trend (- - -) line is indicated.

**Figure 10:**
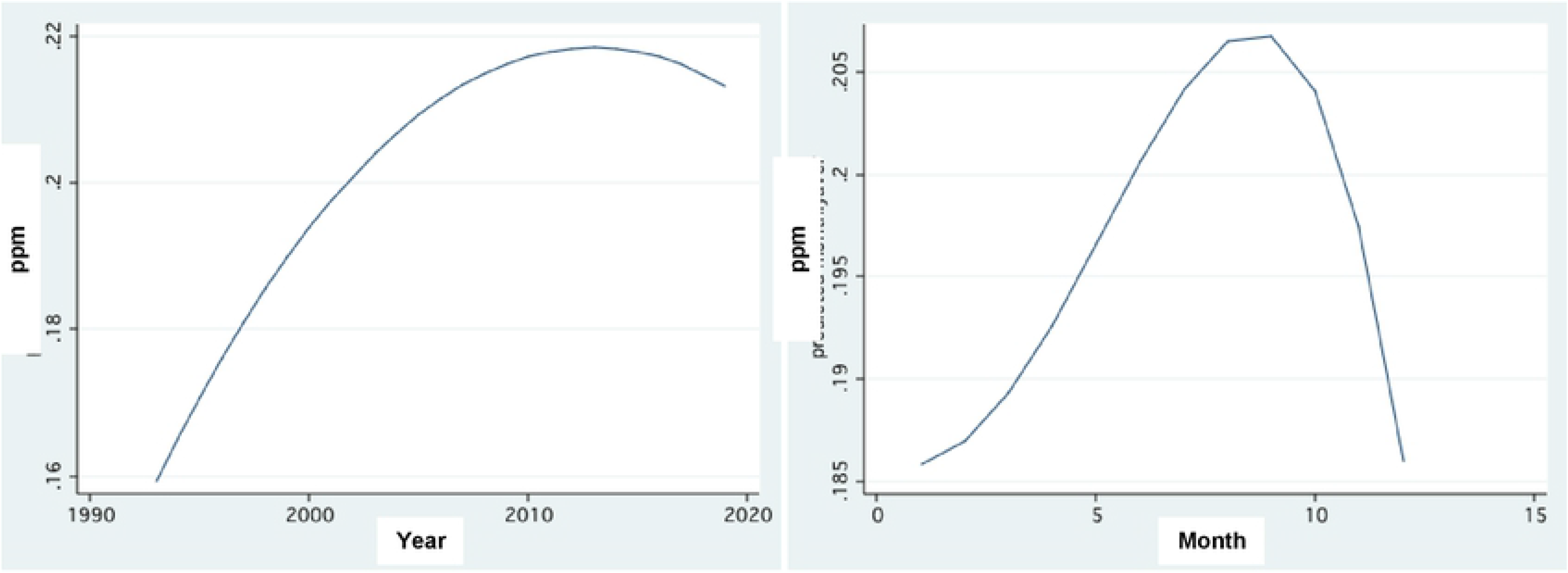
Fractional Polynomial Regression of Temporal changes (annual-left, Monthly - right) of Total Phosphurus averaged concentrations (ppm) of the entire Hula Valley runoff pathways During 1993 - 2020.

**Figure 11:**
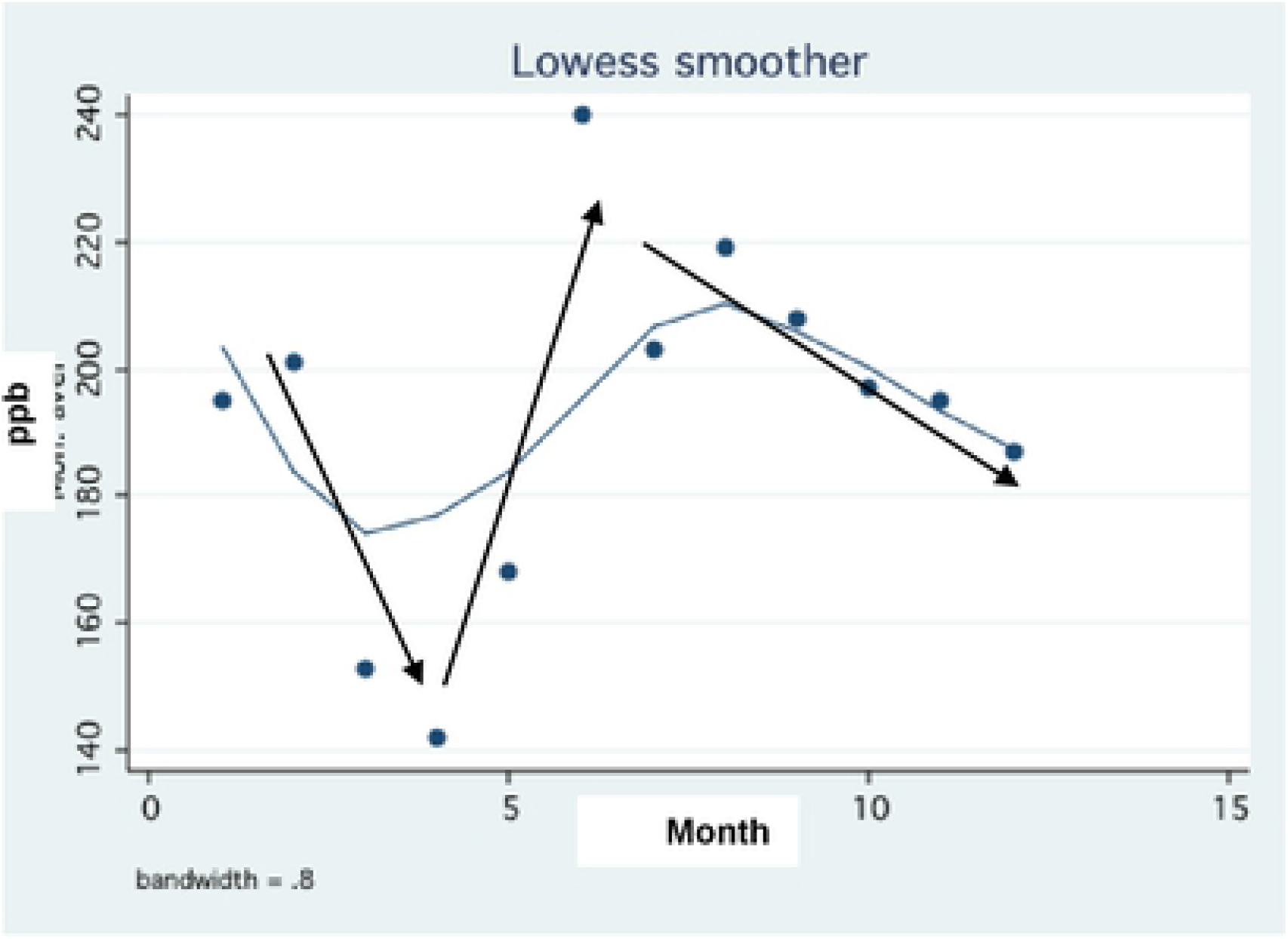
Scatter and Lowess Smoother (Bandwidth 0.8) plot of monthly averages (1993-2020) of TP concentration (ppb) in Agmon-Hula effluent. Solid lines - approximation of seasonal changes.

Critical indication of potential additional P is aimed at both Lake Agmon-Hula ecology (vegetation and phytoplankton) and P flux through Agmon-Hula outflow and partly through River Jordan discharge. The TP mass through Lake Agmon-Hula outflow was found to vary between 0.9–1.6 t/y and the multi-annual mean range of TP concentration in the Agmon-Hula outlet was 0.01–0.2 ppm, and no long-term changes were documented.

The implemented reconstruction of the lost Lake Hula native flora and fauna indicated approximately 300 bird species observed in the Hula Valley. Cranes (Grus grus) are mentioned in this remarkable avifaunal record only twice. Until the early 1990s, only a few cranes visited the Hula Valley. Since then, the valley has been populated annually from November through March by increasing numbers of cranes, up to 50000 in the winter of 2019–2019 (Figure 12). The item that attracts the wintering cranes to the valley is certainly leftover peanut crops. Peanut became an economically successful crop suitable for the heavy organic peat soil in the valley routinely cultured. Efficient agricultural management in the Hula Valley is an essential major objective of the Hula Project. Peanuts are harvested in late autumn and a lot of seeds are left exposed on the ground. The leftover seeds are preferred by the cranes which stay over in the valley while migrating from Europe to Africa. One to two months later, rainfall starts increasing the soil moisture and the leftover seeds begin to ferment. Then the cranes would not like the seeds again and would look for another food source. Consequently, damage is caused to other crops in the Hula valley. Cranes are protected by International Laws and shooting them is not illegal and deportation is possible by other technologies. A collaborative solution between farmers, nature authorities, water managers, land owners, and regional municipalities was budgeted for and implemented. Money was allocated for the rental of a 40-ha field block in the valley to serve as a "feeding station" where purchased corn seeds are given to the cranes twice a day. Feeding starts in late December and continues until early March when the cranes fly back to Europe for breeding. Cranes which arrive before mid-December are partly deported to reduce the number of potential feeders, prevent damage and reduce money spent on corn seeds. This achievement yields benefits for both the landowners and farmers as half a million bird visiting watchers (charged visit), and the Hula Valley effluents are not significantly deteriorated. Moreover, the cranes have been allocated underneath terrestrial eucalyptus trees for where to stay at night, and there they become vulnerable to predators (fox, wolf, mongoose, jackal). Therefore, the bird flocks are beginning to change their night stay location to the protected refuge site in the newly created shallow lake, Agmon-Hula.

**Figure 12:**
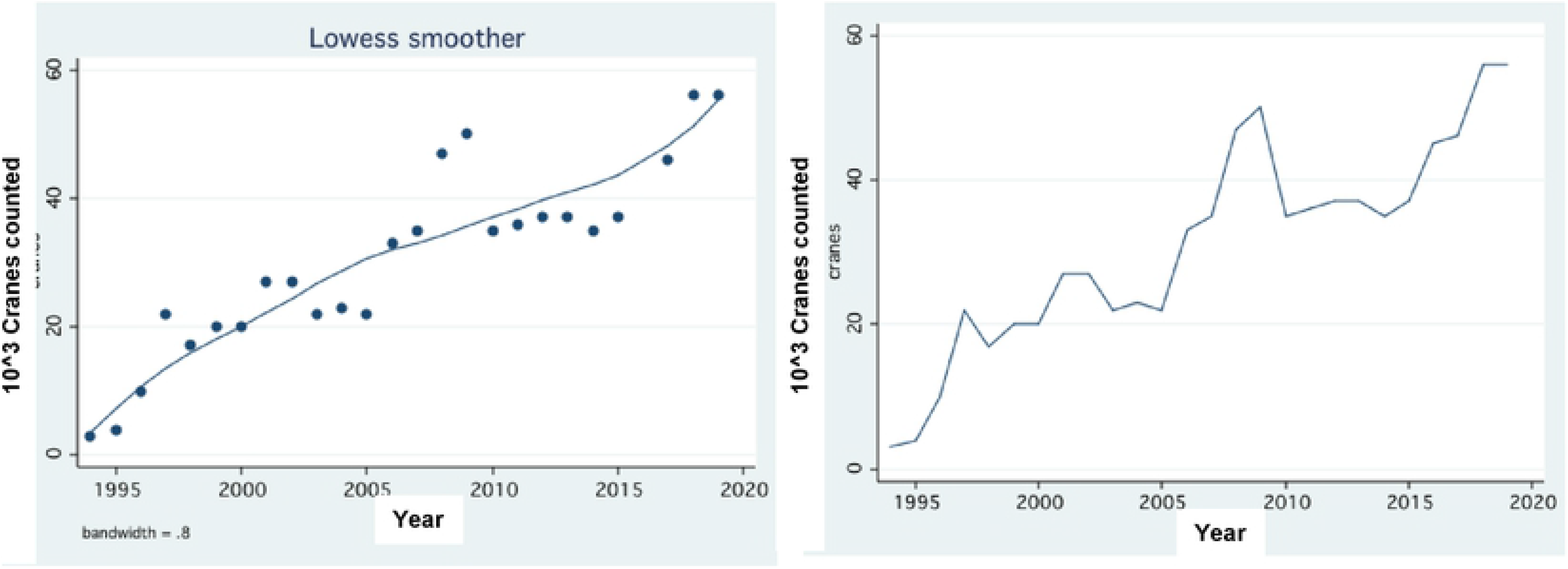
Temparol Crane density (maximal counts) in the Hula Vally during 1995-2019; Left: Lowess Smoother (Bandwidth 0.8); Right: Scatter line plot

The monthly water balance (10^3^ m^3^/season) of Lake Agmon-Hula is exemplified by the 2001 annual summary (Gophen et al. 2003) given in Table 1:

**Table 1:** Seasonal summary of water balance (10^3^ m^3^/season) in Lake Agmon-Hula. Winter – December and January – May; Summer – June – November. Water increment sources are: reconstructed Jordan, Hula East, Canal Z and precipitation; water deficit sources are: outflow and evaporation. Plus shows increment, minus shows deficit. Evaporation (mm/season) and precipitation (mm/season) have been transformed to areal (110 ha) capacity (m^3^3/season).

| Source | Winter | Summer | Annual |
| --- | --- | --- | --- |
| Hula East | 310 | 180 | 490 |
| Reconstructed Jordan | 823 | 967 | 1790 |
| Canal Z | 2080 | 3920 | 6000 |
| Precipitation | 389 | 84 | 473 |
| Evaporation | 905 | 1409 | 2314 |
| Outflow | 2110 | 3380 | 5490 |
| Summary: Increment | +3602 | +5151 | +8753 |
| Deficit | -3015 | -4789 | -7804 |
| Total Balance | +587 | +362 | +949 |

Because the Agmon-Hula Water Level did not changed annually it is suggested that 949 10^3^ m^3^ of water infiltrated through bottom sediments at a rate of about 1 liter per m^2^ per month. Undoubtedly, those infiltrated waters with fluctuating TP concentrations but their fate and allocation are unknown. The multi-annual mean (SD) of TP concentration in the Agmon-Hula effluents is 117 (SD 80) ppb and the Agmon-Hula outflow is 10 × 10^6^ m^3^ the total load is about 1–1.2 ton of TP,. Conclusively, the annual TP output from the Agmon-Hula is 1–1.2 tons of TP, which is mediated by runoff water and about similar load as bottom infiltration. The fate of runoff water-mediated TP is removal into the irrigation system but that of bottom-infiltrated TP is not fully known. The fate of bottom–infiltrated TP has been tentatively suggested to be migration as subterraneanwater water to the Hula-Valley (Gophen et al. 2014).. Lake Agmon-Hula is located topographically at the lowest altitude of the valley., causing a hydraulic gradient from north to south. A Monthly records (1988–2021) of ground water table (GWT) depths in 40 drills and a full-year chemical analysis of ground water samples indicated the following: GWT in the northern part of the valley is higher than in the southern part, the northern soil type is organic whilst the southern soil type is mineral, the northern seasonal amplitude fluctuations in GWT depth are higher, TP concentration in southern ground water is higher. Consequently, it is suggested that migration of ground water-mediated TP takes over the plastic barrier either underneath or beside. The underground accumulation of TP was evidently confirmed as a result of three factors: hydraulic gradient, enhanced erosive impact due to the mineral soil type, which probably has higher free space, and enhanced evapo-transpiration capacity due to the enhanced erosive impact. Nevertheless, how does TP migration occur? what are the steps of the TP cycling? The TP dynamics in the Jordan River and Lake Kinneret does not provide solution to this critical dilemma.

Data given in Table 2 indicates an abrupt elevation of TP content in the Agmon-Hula in summer and fall seasons, which is the result of massive decomposition of submerged vegetation. Aquatic vegetation initiates annually in spring, incorporating phosphorus from bottom sediments. During the degradation of plant mass, phosphorus is transferred into the water in dissolved and particulate forms A documentation (Geyfman 2000) of 64% of TP load input to Jordan waters during winter months (12 and 1–5) and 36% input in summer period has been presented.

**Table 2:**
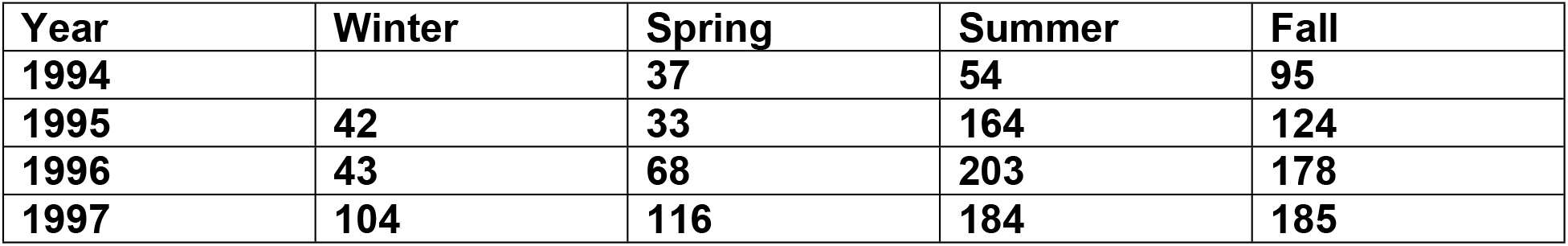
Annual (1994–1997) seasonal averages of total phosphorus concentrations (ppb) in the Lake Agmon-Hula outlet: Winter (January–March), Spring (April–June), Summer (July–September), Fall (October–December).

Data shown in Table 3 indicates the inter-annual dissimilarity of the TP balance due to the respective TP load relation to the water balance; the water balance is dependent on rainfall and agricultural allocation (irrigation) (Table 3).

**Table 3:** Monthly mass balance (kg/month) (input minus output) of TP in Lake Agmon-Hula in 2003 and 2004 (positive value=retained; negative value=deficit) Input sources are: Peat soil drainage through Canal Z and Canal East and the reconstructed Jordan Branch.

| Month | 2003 | 2004 |
| --- | --- | --- |
| 1 | 8 | 180 |
| 2 | 7 | 60 |
| 3 | -63 | 19 |
| 4 | 132 | 10 |
| 5 | 12 | 30 |
| 6 | 79 | 110 |
| 7 | 49 | 70 |
| 8 | 72 | 40 |
| 9 | 23 | 40 |
| 10 | 31 | 10 |
| 11 | -3 | -60 |
| 12 | -210 | 0 |
| Annual Balance | 137 | 600 |

Data shown in Table 4 indicates much higher TP export from Lake Agmon-Hula during drought seasons and, therefore, irrigation intensification, accompanied by TP flushing from the peat soil. Nevertheless, annual TP output from Lake Agmon-Hula ranged between 0.5 and 1.5 tons per year.

**Table 4:** Annual TP mass balance (input minus output: kg/year) in Lake Agmon-Hula (kg/year) for high (2003, 2004) and low precipitation ranges (2007, 2008).

| Year | Annual Balance (kg/y) | Input (kg/y) | Output (kg/y) |
| --- | --- | --- | --- |
| 2003 | 137 | 927 | 792 |
| 2004 | 600 | 1030 | 430 |
| 2007 | -710 | 832 | 1542 |
| 2008 | -620 | 970 | 1590 |

A survey of the distribution and TP content of submerged vegetation in lake Agmon-Hula was carried out for the period 1997–2004. The total dry weight and TP content are given in Table 6.

**Table 6:** Annual averages of total vegetation loads of dry weight and TP (ton) for the entire lake during the period 1997–2004

| Year | Dry weight (t) | TP (t) |
| --- | --- | --- |
| 1997 | 268 | 0.9 |
| 1998 | 213 | 0.7 |
| 1999 | 432 | 0.8 |
| 2000 | 343 | 0.9 |
| 2001 | 740 | 1.2 |
| 2002 | 817 | 1.2 |
| 2003 | 140 | 0.3 |
| 2004 | 698 | 1.1 |

## 4.) Discussion

The impact of climate change during the Anthropocene Era on the ecology of the Lake Kinneret drainage basin (2730 km^2) is widely documented (Figures 1–5) (Gophen 2021). Significant symptoms of these climate changes were, among others, decline in rainfall, reduction in river discharges and, consequently, water-mediated nutrient capacities, temperature elevation, and decline of the ground water table (GWT) in the Hula Valley. Campbell and Capece (1999) documented that P transport mechanism is based primarily on surface topography, and distribution of P between the surface and ground water is related to hydrologic and topographic factors rather than land use intensity. The altitude of the northern part of the Hula Valley is higher than that of the southern section, resulting in a hydrologic gradient from north to south. A wide variety of water-mediated phosphorus resources in the Kinneret watershed are known. One of their principles of these phosphorus resources is agricultural loading (fertilization). Practical fertilization regimes in the Hula Valley is estimated to supply 150–200 ton of P for 20,000 dunam (5–10 gP/m^2) vegetable crops, assuming P fertilization for wheat and corn is negligible. Studies carried out in the Everglades indicated low phosphorous output concentration when the total P mass loading was less than 1 g/m^2^, suggesting that the "one-gram assimilative capacity rule" may be a fairly good approximation for freshwater wetlands (Richardson 1999). The Hula Valley is presently a drained freshwater wetland crossed by join three headwaters into the Jordan River, where the total measured phosphorus content as averaged for 48 years (1970–2018) was 72 tons (Table 4). Consequently, subtraction of effluent capacity (72 t) from loaded masses indicate residual portion of 78–128 t channeled to plant-harvested plant matter, absorbed by soil particles, and migrated into subterranean water pathways, which drained into unknown spaced. Much of the organic P in organic compounds is associated with soil particles, and P transformations are controlled by a combination of P concentrations in solutions and biological activities, of which the most important are microbial alterations of redox reactions and bonding to soil particles (Wetzel 1999). Information about the continuous high loading from commercial fertilizer may contribute significant quantities of soluble P to surface and subsurface drainage (Campbell and Capece 1999). Phosphorus fertilization regimes in the Hula Valley were investigated and criticized by Barnea (2009). The impact of P fertilization rates (4.3–8.6 gP/m^2) on P dynamics in organic and mineral soil types were studied. The low hydraulic conductivity (1 mm/day) and high P adsorption (900–1400 mgP/kg) on organic peat soil which dominates the northern part of the valley significantly reduce underground drainage of P and, probably, further leakage into the environment. On the other hand, hydraulic conductivity of the marl soil in the southern part of the valley is 1.7×10^5^ mm/d, and P migration is higher and mostly occurs as particulate P (Barnea 2009). Therefore, higher P contamination of the environment through underground water mediation is predicted. A flux estimation of 1.0 kgP/ha through free space macropores into the shallow underground water bulk was suggested (Gophen et al. 2014).

The devastated collapse in the abrupt rise and fall of the dense vegetation cover of cattail (Typha domingensis) in the newly created Lake Agmon–Hula wetlands is significantly correlated with the peat soil phosphorus availability (Gophen 2000; Symhayov et al. 2013). The strong correlation between the spatial distribution of cattail and soil P concentration was documented by Miano and DreBusk (1999): soil phosphorus enrichment enhanced macrophyte production and photosynthesis; nutrient storage in plant tissues was correlated to P gradient and cattail rhizome expansion. It was earlier concluded (Gophen 2000) that digging in the Agmon-Hula wetland (“Hula Project” construction, 1993) exposed organic peat soil, enhanced P oxidation and consequent availability, and led to the outbreak of cattail vegetation. Water coverage of the Lake Agmon-Hula bottom due to water-eliminated soil oxidation and P deficiency resulted in the cattail collapse. The consequent future projection is probable enhancement of free available P as a result of decline in soil moisture (Yatom and Rabinovich 1999) resulting acceleration of bounded P release from organic particles. The design of agricultural P fertilizer loading and implementation of attractive environmental systems with reasonable crane-carrying capacity is indispensable. The practical management has to be profitable for both agricultural revenue and eco-touristic activity of bird (crane) watcher visitors despite that the Lake Kinneret water quality needs to be carefully protected. The Hula Valley and Lake Kinneret are twin ecosystems and the flux of pollutants, mostly phosphorus, from the Hula Valley downstream into Lake Kinnert must be controlled. The “Best Management Practices” program that was recommended by Lzuno and Whalen (1999) include categories of fertilizer management, water management and particulate transport reduction. One experimental study (Debusk and Dierberg 1999) demonstrated the potential of vegetation and chemical management to enhance P removal rates in treatment wetlands and effluent P concentration. Application of these recommendations to the Hula–Kinneret ecosystem indicates conclusively that surplus P fertilizer loading and crane droppings presently do not threaten the Kinneret water quality.

Results given in Figure 5 indicates inverse relations between TP concentrations in Agmon-Hula and Jordan waters. This difference is probably due to the dissimilarity between the driving factors which control TP supply to the Agmon-Hula and Jordan waters. In the Agmon-Hula body of water, enhancement of TP is due to the intensive growth rate of aquatic plants in the spring–summer and their decomposition, whilst the rate of discharge controls TP concentration dynamics in Jordan waters. In winter, TP concentration in Agmon-Hula declines but increases in summer, whilst TP increases in Jordan waters during winter when rain and discharge are maximal. Is TP concentration dynamics in Agmon-Hula and River Jordan dependent or independent? This paper suggests it is independent. Deeper evaluation is likely to conclude that phosphorus loading in the Hula Valley is transported into plant matter (harvested crops) and absorbed by soil particles, and the excess migrates into unverified free space. Data shown in Figures 1, 2, 7 and 9 indicate the consequences of the dissimilarities in TP concentrations in Jordan and Agmon-Hula waters to seasonal dynamics. Low TP concentrations in Jordan River waters (Figure 2) are correlated with decline in Jordan discharge. Low level of discharge is typical to summer periods. On the contrary, the TP concentration in the Agmon-Hula waters increases during summer months (Figure 9). These contrary developments confirm the independence linkage trait between Agmon-Hula and River Jordan bodies of water. Moreover, the linear regression between rainfall regime (obviously in winter) and Jordan discharge (Figure 1) was found to be significant (r^2^ = 0.3268, p = 0.0044). Figures 8, 9 (left panel), 10, 13, and 14 indicate temporal elevation of TP concentration in Lake Agmon-Hula between 1994–2020. The opposite temporal changes in nutrient concentrations in the Jordan waters are shown in Figures 11 and 12. Nevertheless, Figures 11 and 12 probably confirm that the presence of cranes in the Hula Valley had no impact on nutrient concentrations in the Jordan waters. The precautious trait of crane as phosphorus contaminators became realistic as a result of their daily migration to the Lake Agmon-Hula for night stay as a way to protect themselves from predators. Surprisingly, the TP concentration in the Agmon-Hula waters was found to lower in winter when the cranes were present and to increase in summer several months after the cranes’ deportation. The increase in TP in Agmon is therefore evidently due to the submerged vegetation growth and degradation. The difference between TP point measured concentration and phosphorus mass balance needs to be clearly stated (Table 4). The P mass balance (total input minus total output) in Lake Agmon during two drought years of 2007 and 2008 was negative, i.e. much of the input as well as plant-mediated P was retained, but positive in wet seasons. The residence time (RT) of lake water is shorter (wet season) when more P is flushed out. In dry seasons RT shortens and water exchange level declines when Phophsphorus mass is retained.. Two environmental ecosystems have an impact on P concentration in the Agmon-Hula waters: 1) the Lake itself with submerged plants and chemo-physical processes (sedimentation, phosphorus cycling, etc.) (Gophen 2000; Gophen et al. 2001; Symhayov et al 2013); and 2) peat organic soil and water (precipitation, irrigation) flushing (Yatom and Rabinovich 1999; Yatom et al. 1996; Litaor et al. 2013, 2014; Haygarth et al. 2013; Reichman et al. 2013). Plant-mediated phosphorus and geo-chemical processes in the peat soil have a major impact on the lake ecosystem, with the first dominant in summer and the second in winter.Considering that plant-mediated phosphorus has a major impact on the lake ecosystem and geo-chemical processes inside the Peat soil as dominant, the domination of the first occur in summer and that of the second in winter. Plant-mediated phosphorus probably contributes the highest amount of phosphorus to the external environment.

**Figure 13:**
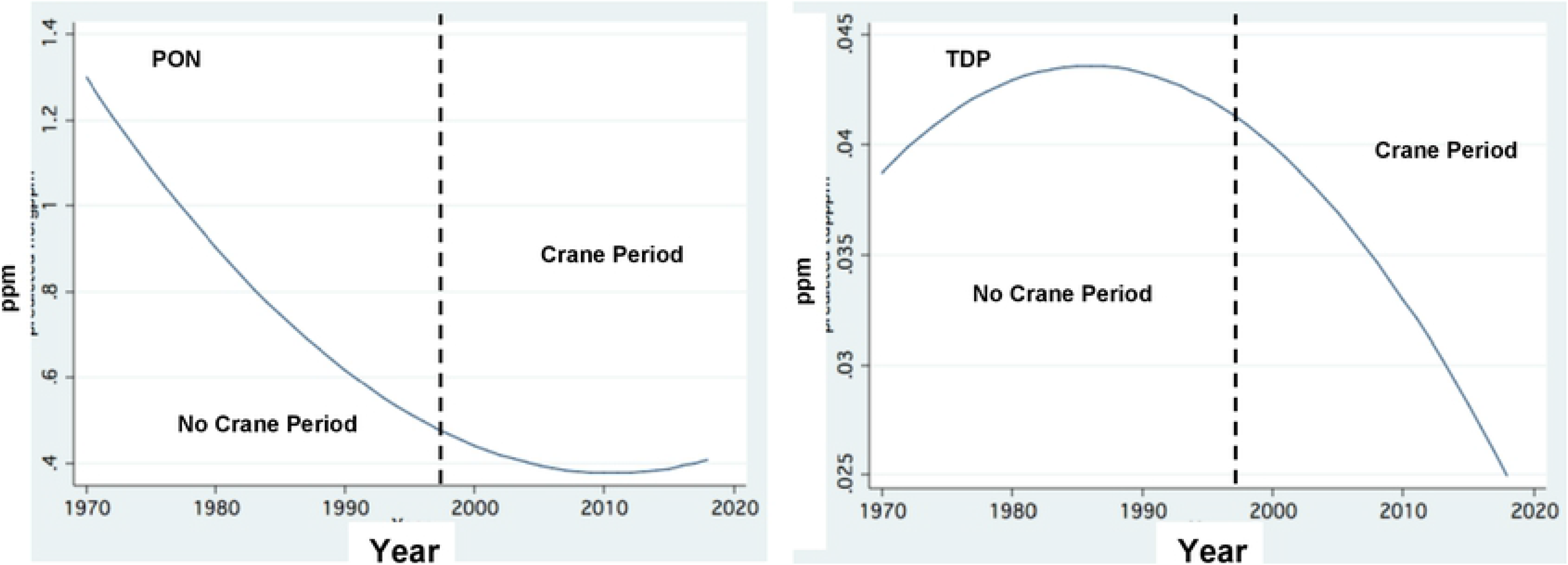
Fractional Polynomial Regression of annual means of temparol (1970-2018) changes of The concentrations (ppm) of Total Dissolved Phosphorus (TDP) (right) and Particulate Organic Nitrogen (PON) (left) in the Jordan Waters. Broken line limits between with and without Cranes in the Valley.

**Figure 14:**
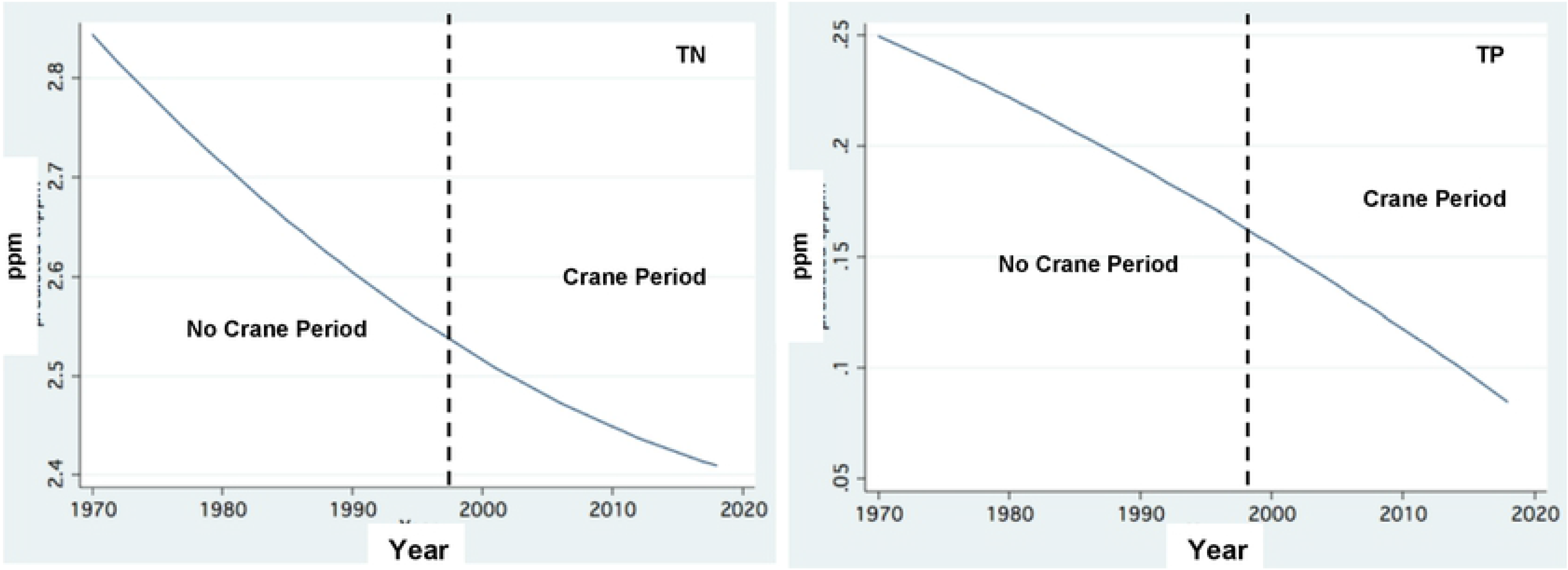
Fractional Polynomial Regression of annual means of temporal (1970-2018) changes of The concentrations (ppm) of Total Phosphorus (TP) (right) and Total Nitrogen (TN) (left). Broken line limits between with and without Cranes in the Valley.

Although the impact of geo-chemical processes in organic peat soil is significant, Yatom et al. (1996) found that wet-dryness processes in peat soil are significant as well, since P is thus realized and transported by water. Results shown here probably indicate low rates of P leakage from the high-loaded P fertilizer (5–10 gP/m^2^) such that the peat soil substrate has a loiading capacity that is below maximum (Richardson 1999). Therefore, P contribution to the environment, although supplemented by crane droppings, is presently not risky.

It was clearly indicated that the increase in TP concentration in the Agmon-Hula waters occurs in summer month, mostly resulting from degradation of the biomass of emergent, submerged, and floating high plants (Phragmites spp., Potamogeton spp., Najas spp., Myriophyllum sp.) and sediment mat cover comprising of algal vegetation biomass. The concentrations of plant-mediated TP and SRP (Soluble Reactive Phosphorus) in the Lake Agmon-Hula waters during summer months (Gophen 2000) were higher than in the P input sources (Canal Z and reconstructed Jordan). It was estimated (Gophen 2000) that the degradation of aquatic plant biomass, in addition to the external input, contributed approximately 325 kgP to the total load in the summer of 1999. One experimental study (Markel 1998) documented very low rates of advection (upward) flows through the bottom sediments, contributing about 7 ppb of P into the thin and anoxic cover layer in 24 hours. The foundation of bottom anoxic layer is not permanent and alternatively a concentration of 7–8 µM of FeS was documented (Markel 1998). Kaplan (1998) documented 268 tons of dry matter of aquatic plants containing 1 ton of phosphorus. How much of this P load transferred into the water or settled at the bottom was not measured but plant-mediated P sourcing was confirmed. Hydrological conduction in the Agmon-Hula Lake system where residence time is short enough was found to be suitable to achieve optimal maintenance of agricultural and eco-touristic objectives together with protection of the Kinneret water quality. The impact of water exchange control by inflow–outflow regimes is critical for the prevention of P elevation and eutrophication in lakes, reservoirs and wetlands (Volohonsky et al 1992). Nevertheless, the newly created Agmon wetlands ecosystem could not prevent the abrupt outbreak of dense Typha vegetation immediately after filling water in the Lake Agmon-Hula. We confirmed that the reason was the short-term surplus availability of P. Shortly after, P availability reduced dramatically and the Typha vegetation waned (Miao and DeBusk 1999). Later on, as a consequence of inundation due to water level fluctuation, P availability along the narrow beach stripe increased and Typha vegetation was renewed. The organic peat soil in the Hula Valley is a P-rich habitat (Litaor et al. 2013, 2014; Reichman et al. 2013; Haygarth et al. 2013). Nevertheless, there is a storage of 22–37 and 11–22 tons of P in the upper two 5-cm layers, respectively, in the Agmon-Hula bottom sediments (Gophen 2000). Root system penetration of Typha probably enables efficient utilization of P in the upper and lower layers, making the available P stock larger. Long-term study of the Lake Agmon-Hula region confirmed Agmon-Hula system as a phosphorus contributor to the runoffs in the vicinity (Gophen 2015a, b) but probably not further. Nutrient inputs into Lake Kinneret are probably not significantly influenced. TP concentrations in Jordan River are rather stable and not significantly affected by Agmon. Erosion eco-forces produced by headwater river discharges are suggested to have a significant impact on the water-mediated P-carrying capacity. An 11-year (1994–2004) record indicated mean concentration of TP in Canal Z as 0.11 ppm, in the Agmon-Hula outlet as 0.15 ppm, in the reconstructed Jordan as 0.11 ppm, and in Hula east as 0.2 ppm. Moreover, linear regression confirmed that in Canal Z and in the Agmon-Hula outlet, summer TP concentrations are significantly higher than the winter values (Gophen 2015a, b).

The massive wintering of cranes in the Hula Valley started in the early 1990s. Cranes usually stay during spring–summer months in European territories to breed and take care of their young. Migration of cranes to the south happens naturally during fall via two major routes: western towards Spain and eastern towards East Africa through Israel. Until the early 1990s, the number of crane landings in winter reached a maximum of a few thousands allocated sporadically in northern parts of Israel. Nevertheless, as a means of agricultural development in the Hula Valley, peanut cultivation was begun; the peanuts are harvested in fall, leaving plenty of residual seeds on the ground surface. These leftover peanut seeds were discovered by the migrator cranes and their winter landing for feeding was routinely initiated. Nevertheless, rainfall wetting caused fermentation of the peanuts; the fermented peanuts became unpalatable for the cranes and in their search for other food sources they caused damage to agricultural crops. When the rate of landing and crop damage increased, the need to find a solution became inevitable. Thus, purchased corn seeds were scattered for the cranes in an uncultivated field block. This sophisticated solution initiated difficulties: the corn seeds were favored by the cranes but were costly and their consumption rate was high, resulting in higher landing rates. When they were no longer supplied the cranes reverted to damaging crops with renewed intensity.the corn seeds were favored by the cranes but were costly and their consumption rate was high, resulting in higher landing rates and intensified crop damage. The most problematic issue came when the cranes began to utilize Lake Agmon-Hula as night shelter for protection from natural predators (fox, mongoose, wolf, coyote). The complexity of the management parameters initiated a risky structure: Increase in the number of external bird flocks led to higher consumption of imported food. and contributed nutrients either to terrestrial land or directly into the Lake Agmon-Hula waters. Crane droppings contributed about 5.24 gP/individual/day (there were about 50,000 cranes), making a total daily loading of 262 ton of phosphorus in the Hula Valley, with approximately 50% or more ending up directly in the Lake Agmon-Hula waters (Gophen 2017). What is the fate of such an intensive P loading? Results (Figure 7, 8, 9) clearly indicate that there is less decline in TP concentration in the Agmon-Hula water when the cranes stay in the Valley. There was a temporal decline in nutrient concentrations (particulate organic nitrogen, total dissolved phosphorus, total nitrogen, total phosphorus) (Figures 11, 12) between 1970 and 2018. Continuous decline in Phosphorus from 1970, before the accumulation of the migrator cranes in winter, was not interrupted by the wintering stay 20,000–50,000 cranes fed on corn seeds. During winter months, submerged vegetation is negligible. Therefore, water-mediated P effluents, bottom infiltration and particulate sedimentation as potential removal channels are suggested. Nevertheless, no indication of enhanced P input into Lake Kinneret through Jordan discharge was documented. Conclusively, it can be suggested that the massive damage caused by the Cranes Crane massive practical damage (not through nutrient enrichment) is critical to agricultural crops. The time table for corn seed supply is under managers’ control. Therefore, before feeding starts, aggressive deportation is implemented to reduce the crane population: in 2020, the maximum number of cranes was 33000 whilst a year earlier it was 56000. Moreover, the Kinneret and Lake Agmon-Hula water qualities are not endangered. Linear prediction of annual averages of Jordan water-mediated TP input (ton/y) and the annual discharges indicated, obviously, significant correlation (r^2 = 0.596, p < 0.0001). The long-term decline in the Jordan discharge from 15 to <10m^3^/s since the mid-1980s is due to climate change. Moreover, since the Hula Reclamation Project was accomplished beside bird watching (thousands of cranes and another 175 documented species) attraction also the agricultural revenue was doubled (Znovar et al 2010). Conclusively, precautionary concerns due to the Jordan and Lake Agmon-Hula water-mediated P are not presently confirmed. Optimization of crane watching attraction should prevent agricultural damage due to effective deportation of cranes in fall and corn seed limitation in dedicated field block (feeding dedicated plots of land, “Crane 5-star Restaurant”). The Eco-Touristic Crane Project was designed to be a part of a comprehensive objective aimed at enhancement of ecosystem sustainability. The solution can be conclusively summarized thus: to reduce agricultural damage by feeding the crane seeds with corn seeds on the same land, where the crane birds often gathered during the day time, and left this area for night stay in the shallow lake where they felt protected from predators. Bird watchers visit, and the management of the Hula project removes nutrients from the Kinneret loads. This crane project represents an efficient marrying of touristic bird watching and limnological interests to prevent eutrophication in Lake Kinneret.This Crane project represents an efficient partnership of coexisting birds and limnological interests for the prevention of Eutrophication in Lake Kinneret.

The Hula Reclamation Project was aimed at ensuring sustainability of modified eco-systems by agricultural development, Kinneret water quality protection and nature conservation. The tension between farmers, water managers, and nature preservers was reduced, leading to their collaboration instead. The outcome of the Hula Project was ecosystem renewal, leading to the development of a tourist attraction and enriching the biological diversity with approximately 300 species of birds, including 40,000–56,000 wintering Cranes annually, 40 species of water plants, and 12 species of fish. The new ecosystem of the shallow Lake Agmon-Hula with the surrounding Safari habitat ecosystem became a tourism attraction. Potential resource contributors to water-mediated phosphorus include the following: Kinneret headwaters (Table 5), Lake Agmon-Hula through crane droppings, aquatic vegetation, and the major peat soil-drained water pathways (Table 1) in the Hula Valley. The Kinneret watershed region has undergone changes in climate condition of which dryness is emphasized. These changes enhanced processes of decline in rainfall–river discharge, accompanied by changes in nutrient dynamics and reductions in input concentrations (particulate organic nitrogen, total dissolved phosphorus, total nitrogen and total phosphorus) (Figure 13,14). It is likely that these modifications in nutrient dynamics were not affected by the presence of cranes in the Hula Valley, even though the seasonal changes in TP concentration in the Agmon-Hula effluent are due to the onset and offset of submerged macrophytes (Figure 15). The status of the phosphorus cycle in the Hula Valley is not clearly known.. Therefore, a tentative conclusion can be stated as follows: Phosphorus input into Lakes Agmon-Hula and Kinneret is affected mostly by climate change and submerged macrophytes and, to a lesser extent (if at all), by cranes.

**Table 5:** Total annual phosphorus loads in three headwaters—Hatzbany, Banias, and Dan—and from Hula Valley as averaged for 48 years (1970-2018) (% role is indicated).

| Source | Ton P/y |
| --- | --- |
| Hatzbany | 11.5 (16%) |
| Banias | 10.8 (15%) |
| Dan | 5 (4.2) |
| Hula | 46.1 (64%) |
| Total | 72 |

**Figure 15:**
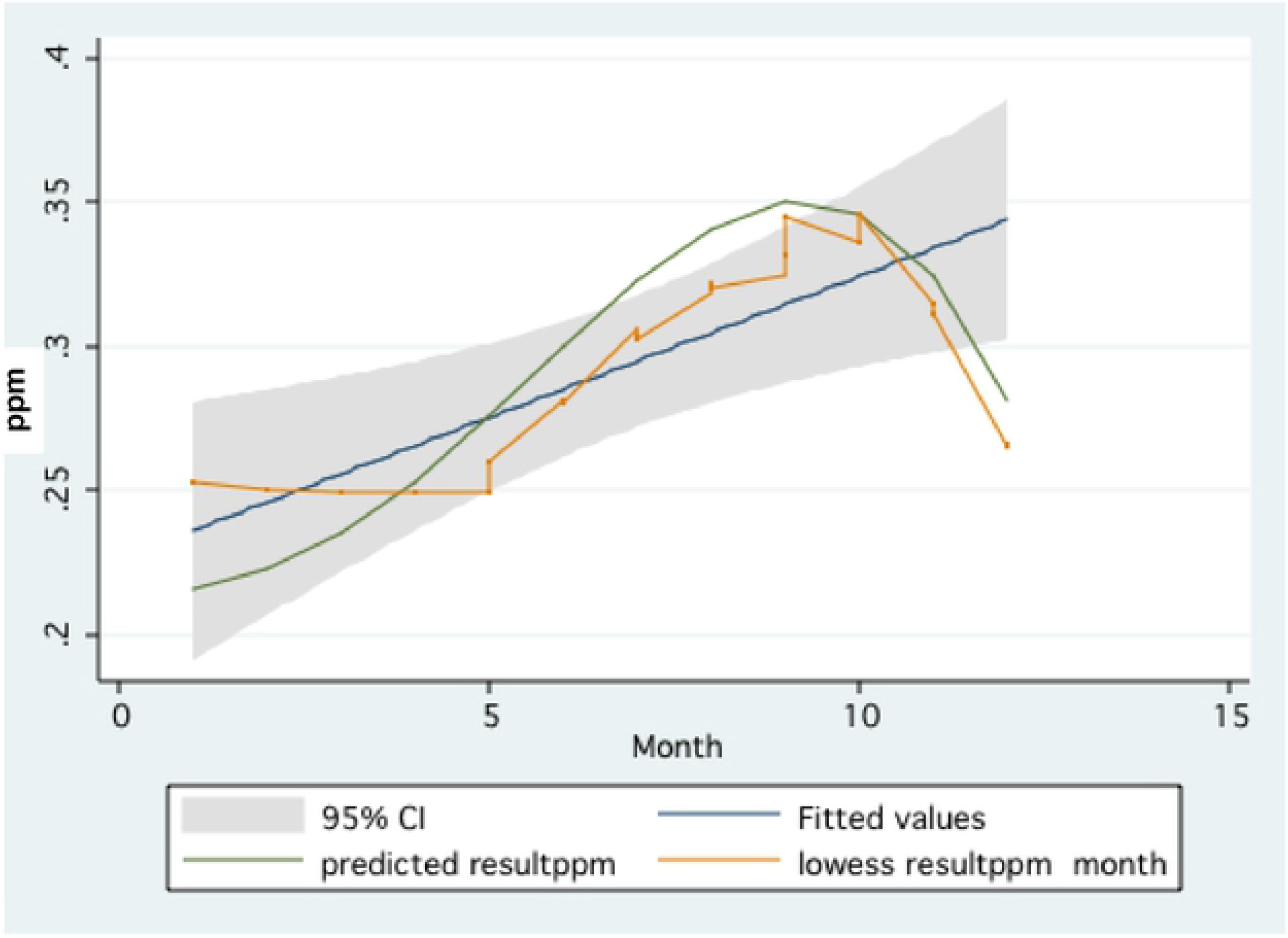
Three statistical method expressions of seasonal (1-12 months) changes (averaged for 1993-2020) of TP concentrations (ppm) in the Lake Agmon-Hula effluents: 1)Linear Prediction of Fitted value with 95% CI; 2) Predicted Value by Fractional Polynomial; 3) LOWESS (0.8).

## 5.) Conclusive remarks

Sources of phosphorous supply to Lake Kinneret within the drainage basin that are discussed in this paper are: natural soil and rock beds eroded by headwater discharges, components of organic peat and marl soil in the Hula Valley, winter migratory crane droppings, and external P fertilizer loading. Two other sources, which are not considered, are dust deposition and Kinneret internal bottom (chemically–microbiologically released). External fertilizer loading is partly incorporated into harvested crops and adsorbed on soil particles. Nevertheless, the fate of the rest of the P supply, which migrates mostly in particulate form, is unknown, but probably not end up and fully included in the Jordan River discharge. sIt is suggested that the present input regime of phosphorus into Lake Kinneret is not risky but the eco-hydrological structure of the ecosystem has a reasonable potential which may lead to eutrophication in both Lake Kinneret and Lake Agmon-Hula. It was indicated that 77% and 68% of total TP load in River Jordan and its headwaters respectively were fluxed during winter months (October–April). It can therefore be concluded that this difference resulted from the higher hydraulic erosion effect maintained by the winter discharges. The impact of agricultural activities and cranes is minor, even though the difference between winter and summer TP loads in the Hula Valley`s effluents is pronounced (higher in summer). Conclusively, the dominant factor that controls water-mediated TP concentration and, consequently, load of the Kinneret inputs is erosion produced by friction. The higher the discharge, the higher the TP concentration. Additional resources such as migratory birds, external fertilizer loads, and, probably to a lesser extent, dust deposition are balanced within the ecosystem compartments, and the fate of the surplus P is unknown.

## Reference

Barnea, I.(ed) 2008, Hula Project Annual Report, Jewish National Fund (Keren Kayeemet LeUsrael) Migal-Scientific Research Instoiitute and Israeli Watr Authority159,p.

Barnea I,(d) 2008-2018, Hula Project Annual Report, Jewish National Fund (Keren Kayeemet LeUsrael) Migal-Scientific Research Instoiitute and Israeli Watr Authority 232p.

Barnea, I. 2009, Reexamination of Phosphorus Fertilization Practices in the Altered Wetland Soil of Hula Valley, Israel. Thesis: Master of Science, Faculty of Agriculture, Food and Environmental Quality, The Hebrew University,Jerusalem; 104 p.

Campbell, Kenneth L., and John C. Capece., 1999, Chapter: Hydrologic Processes Influencing Phosphorus Transport, in: Reddy, K. R., G. A. O’Connor and C. L. Schelske (eds.) 1999, Phosphorus Biogeochemistry in Subtropical Ecosystems, LEWIS PUBLISHERS Boca Raton London New York Wshington D.C. pp 343–354.

DeBusk, Thomas A., and Forrest E. Dierberg 1999, Chapter:Techniques for Optimizing Phosphorus Removl In Treatment Wetlands; in: Reddy, K. R., G. A. O’Connor and C. L. Schelske (eds.) 1999, Phosphorus Biogeochemistry in Subtropical Ecosystems, LEWIS PUBLISHERS Boca Raton London New York Wshington D.C. pp 467–488.

Foner, H. A., E. Ganor, and G. Gravenhorst, 2009, The Chemical composition and sources of the bulk deposition on Lake Kinneret (The Sea of Galilee), Israel, Journal of Arid Environments, Volume 73, Issue 1, January 2009, Pages 40–47; https://doi.org/10.1016/j.jaridenv.2008.09.013

Geyfman, Y. 2000, Multi Dimensional Analysis of the Dynamics of Jordan Loads to Lake Kinneret during 1970-1999 Unsupervised Neural Networks (Supplements) (in Hebrew). 16 p.

Gophen, M., 2000 a. The Hula Project: N and P dynamics in Lake Agmon and pollutants removal from the Kinneret inputs.Water Science and Technology Vol. 42 No. 1-2 pp 117–122.

Gophen, M., 2000, Nutrient and plant dynamics in Lake Agmon Wetlands (Hula Valley, Israel): a review with emphasis on Typha domingensis (1994-1999). Hydrobiologia 00: 1–12.

Gophen, M., D. Kaplan, Y. Tsipris, M. Meron,& I. Bar-Ilan. 2001.The newly constructed wetland ecosystem of Lake Agmon (Hula Valley, Israel): Functional perspectives. In: Treatment Wetlands for Water Quality Improvement; Quebec 2000 Conference Proceedings (selected papers).(compiled by: J. Pries, CH2M Hill Canada Ltd.pp.53–62.

Gophen, M. Y. Tsipris, M. Meron and I. Bar – Ilan, 2003.. The management of Lake Agmon Wetlands (Hula Valley, Israel). Shallow Lakes Conference, Lake Balaton, Hungary, June 2002. Hydrobiologia, 506 (1): 803–809.

Gophen, M., M. Meron, V. Orlov-Levin, & Y. Tsipris, (2014),Seasonal and spatial distribution of N & P substances in the Hula Valley (Israel) subterranean. Open Journal of Modern Hydrology; , 4, 121–131. http://dx.doi.org./10.4236/ojmh.2014.44012.

Gophen, M. 2015 a. Management Improvement of the Agmon Wetlands System (Hula Valley, Israel) aimed at Enhancement of Bird Populations and Kinneret Protection. Open Journal of Modern Hydrology,2015,5, 1–9.

Gophen, M. 2015 b. Nitrogen and Phosphorus dynamics in the Shallow Lake Agmon (Hula Valley, Israel), Open Journal of Ecology, 2015, 5, 55–65; http://dx.doi.org./10.4236/ojmh.2015.51001.

Gophen, M. 2017. Partnerships between the Managements of Cranes (Grus grus) and Kinneret Water Quality Protection In the Hula Valley, Israel. Open Journal of Modern Hydrology, Open Journal Of Modern Hydrology 7, 200–208, .http://doi.org/10.4236/ojmh.2017.72011.

Gophen, M. 2021, Climate Change – Enhanced, Cyanobacteria Domination in Lake Kinneret: A Retrospective Overview; Water (MDPI) 2021, 13, 163, pp. 1 – 19; https://doi.org/10.3390/qw13020163. In: Water (MDPI) Special issue “Cyanobacteria Threat on Freshwater safety”, (Prof. Moshe Gophen, Guest Editor.).

Gophen, M, M. Meron, Y. Tsipris, V. Orlov-Levine, and M. Peres.. 2016. Chemical, Hydrological and Climatological Properties of Lake Agmon, Hula Valley (Israel) (1994-2006). Open Journal of Modern Hydrology, 2016, 6, 8–18. http://dx.doi.org./10.4236/ojmh.2016.61002.

Gophen, M.,and D. Levanon,.(eds) 1993-2006 Hula Project, Annual Reports: Migal-Sientific Research Institute, Jewish National Fund (Keren Kayemet LeIsrael), US Forestry Service International Project, Israeli Water Authority.

Gonen, E.(ed) 2007. Hula Project Annual Report, Jewish National Fund (Keren Kayeemet LeUsrael) Migal-Scientific Research Instoiitute and Israeli Watr Authority, 133 p.

Gvirtzman, H. 2002, Israel Water Resources: Chpters in Hydrology band Environmental Sciences. Yad Ben-Zvi Press, Jewrusalem,287 p.

Haygarth, P. M., A. Delgado; W. J. Chardon; M. I. Litaor; F. Gil-Sotres; and J. Torrent, 2013, Phosphorus in soils and its transfer to water: From fine-scale soil processes to models and solutions in landscapes and catchments Soil Use and Management 2013 Volume 29 Issue SUPPL.1 Pages 03–21.

Kaplan, D. 1998, Chapter: Submerged Vegetation, in: Hula Project Annual Report (M. Gophen ed.)pp. 85–96.

Karmon, Y. 1956,The Northern Huleh Valley: Its Natural and Cultural Landscape. The Magnes Press, Hebrew University, Jerusalem,1956. 108 p. Hadassah Apprentice School pof Printing, Jerusalem, (in Hebrew).

LKDB-IOLR 1970-2018. Annual Reports, Kinneret Limnoilogical Laboratory, IOLR.

Litaor, M.I., I. Chashmonai; I. Barnea; O. Reichmann; M. Shenker 2013;Assessment of phosphorus fertilizer practices in altered wetland soils using uncertainty analysis; Soil Use and Management 2013 Volume 29 Issue SUPPL.1 Pages 55–63

Litaor, M.I. O. Reichmann; E. Dente; A. Naftaly; M. Shenker, 2014 The impact of ornithogenic inputs on phosphorous transport from altered wetland soils to waterways in East Mediterranean ecosystem Science of the Total Environment 2014 Pages 36–42

Lzuno, Forrest T. and Paul J. Whalen 1999. Chapter: Phosphorus Managemnt in Organic (Hitosols) Soils, in: Reddy, K. R., G. A. O’Connor and C. L. Schelske (eds.) 1999, Phosphorus Biogeochemistry in Subtropical Ecosystems, LEWIS PUBLISHERS Boca Raton London New York Wshington D.C. pp 425–445.

Markel, D. E. Sas, B. Lazar, and A. Bein, 1998, Biogeochemical evolution of sulfur-iron rich aquatic system in a reflooded wetlands environment (Lake Agmon, northern Israel), Wetlands Ecology and Management, 6:103–120. 1998 Kluwer Axcademic Publisher, Printed in the Netherlands.Special Issue Destruction and Creation of Wetland Ecosystem in Northern Israel, Guest Editor K.D. Hambright

Miao, L. S.,and W. F. DeBusk, 1999. Chapter: Effects of Phosphorus Enrichment o9n Structure and Function of Sawgrass and Cattail Communities in the Everglades; in: Reddy, K. R., G. A. O’Connor and C. L. Schelske (eds.) 1999, Phosphorus Biogeochemistry in Subtropical Ecosystems, LEWIS PUBLISHERS Boca Raton London New York Wshington D.C. pp 275–299.

Reddy, R., G. A. O’Connor, and C.L. Schelske (eds.) 1999, Phosphorus Biogeochemistry in Tropical Ecosystems; LEWIS PUBLISHERS, Boca Raton London New York Wshington, D.C. 707 p.

Reichmann, O., Y. Chen, and M. Iggy Litaor. 2013, Spatial Model Assessment of P Transport from Soils to Waterways in an Eastern Mediterranean Watershed; Water 2013 Volume 5 Pages 262–279.

Reichman, O. Chen, Y. and M. I. Litaor. 2016,The Impact of Rainfall-Runoff Events on the Water Quality of the Upper Catchment of the Jordan River, Israel. In: Integrated Water Resources Management: Concept, Research and Implementation,2016, Borchardt et al Ed.,Springer NY, Pages 129–146

Richardson, Curtis.J., 1999.The role ofwetrlands in storage, and cycling of Phosphorus on the landscape: A 25 – Year retrospective. Chapter in: Reddy, K. R., G. A. O’Connor and C. L. Schelske (eds.) 1999, Phosphorus Biogeochemistry in Subtropical Ecosystems, LEWIS PUBLISHERS Boca Raton London New York Wshington D.C. pp 47–68.

Simhayov, R.,; M. I. Litaor; I. Barnea; M. Shenker 2013;Catastrophic dieback of Cyperus papyrus in response to geochemical changes in an East Mediterranean altered wetland; Wetlands 2013 Volume 33 Issue 4 Pages 747–758

Volohonsky, H., G. Shaham and M. Gophen. 1992. The Impact of Water Inflow Reduction on Trophic Status of Lakes. Ecological Modelling 62:135–147.

Wetzel, Robert G. 1999, Chapter: Organic Phosphorus Mineralization in Soil and Sediments, Chapter in: Reddy, K. R., G. A. O’Connor and C. L. Schelske (eds.) 1999, Phosphorus Biogeochemistry in Subtropical Ecosystems, LEWIS PUBLISHERS Boca Raton London New York Wshington D.C. pp 225–245.

Xiang, H. F. and A. Banin, 1996. Solid-Phase Manganese Fractionation Changes in Saturated Arid Zone Soil Pathways and Kinetics; American Journal of Soil Sciences (60), pp 1072–1080.

Yatom, S. M. Meron, O. Rabinovich, E. Yasur and A. Banin. 1996. Chapter: Manganese Deficiency and Reduction of Organic Soil Productivity in the Hula Valley, in: 1995 Hula Project Annual Report (Gophen M. editor); pp 8–21. (in Hebrew)..

Yatom, S. and O. Rabinovich, 1999.Chapter, in: 1998 Hula Project Annual Report (Gophen, M. editor): Fractionation of organic soils in the Hula Valley: Partition Indices and Organic Fractionation: Conclusive Report; pp 86–99 (in Hebrew).

Znovar Oved Gobi Ltd.: Shacham, G, H. Tsaban, Y. Avnimelech, and A. Ofer, 2011, Hula Project 2^nd^ Stage, Development Program, Chapter: Opinion about Agricultural, Water consumption, Environmental and Touristic changes in the Hula Valley. Interim Report 31 p. (in Hebrew).

